# Oomycete small RNAs invade the plant RNA-induced silencing complex for virulence

**DOI:** 10.1101/689190

**Authors:** Florian Dunker, Adriana Trutzenberg, Jan Samuel Rothenpieler, Sarah Kuhn, Reinhard Pröls, Tom Schreiber, Alain Tissier, Ralph Hückelhoven, Arne Weiberg

## Abstract

Fungal small RNAs (sRNAs) hijack the plant RNA silencing pathway to manipulate host gene expression, named cross-kingdom RNA interference (ckRNAi). It is currently unknown how conserved and significant ckRNAi is for microbial virulence. Here, we found for the first time that sRNAs of a pathogen representing the oomycete kingdom invade the host plant’s Argonaute (AGO)/RNA-induced silencing complex. To demonstrate the functionality of the plant-invading oomycete *Hyaloperonospora arabidopsidis* sRNAs (*Hpa*sRNAs), we designed a novel CRISPR endoribonuclease Csy4/GUS repressor reporter to visualize *in situ* pathogen-induced target suppression in *Arabidopsis thaliana* host plant. By using 5’ RACE-PCR we demonstrated *Hpa*sRNAs-directed cleavage of plant mRNAs. The significant role of *Hpa*sRNAs together with *At*AGO1 in virulence was demonstrated by plant *atago1* mutants and by transgenic *Arabidopsis* expressing a target mimic to block *Hpa*sRNAs, that both exhibited enhanced resistance. Individual *Hpa*sRNA plant targets contributed to host immunity, as *Arabidopsis* gene knockout or *Hpa*sRNA-resistant gene versions exhibited quantitative enhanced or reduced susceptibility, respectively. Together with previous reports, we found that ckRNAi is conserved among oomycete and fungal pathogens.

## Introduction

Plant small RNAs (sRNAs) regulate gene expression via the Argonaute (AGO)/RNA-induced silencing complex (RISC), which is crucial for tissue development, stress physiology and activating immunity (Chen, 2009; Huang et al., 2016; Khraiwesh et al., 2012). The fungal plant pathogen *Botrytis cinerea*, secretes sRNAs that hijacks the plant AGO/RISC in *Arabidopsis*, and *B. cinerea* sRNAs induce host gene silencing to support virulence (Weiberg et al., 2013), a mechanism known as cross-kingdom RNA interference (ckRNAi) (Weiberg et al., 2015). In fungal-plant interactions, ckRNAi is bidirectional, as plant-originated sRNAs are secreted into fungal pathogens to trigger gene silencing of virulence genes (Cai et al., 2018; Zhang et al., 2016). It is currently not known, how important ckRNAi is for pathogen virulence in general and whether other kingdoms of microbial pathogens, such as oomycetes, transfer sRNAs into hosts to support virulence.

Oomycetes comprise some of the most notorious plant pathogens and belong to the eukaryotic phylum Stramenopiles, which is phylogenetically distant from animals, plants and fungi (Kamoun et al., 2015). We here demonstrate that the downy mildew causing oomycete *Hyaloperonospora arabidopsidis* transfers sRNAs into the host plant *Arabidopsis thaliana* AGO1/RISC, which are functional to silence host target genes, and that invasive oomycete sRNAs are crucial for virulence by silencing plant host defence genes.

## Results

### Oomycete sRNAs invade the plant AGO1

We used the oomycete *Hyaloperonospora arabidopsidis* isolate Noco2 as an inoculum that is virulent on the *A. thaliana* ecotype Col-0 (Knoth et al., 2007). We presumed that *H. arabidopsidis* produced sRNAs, as sRNA biogenesis core components RNA-dependent RNA polymerase and Dicer-like protein were found in the genome (Bollmann et al., 2016). In order to identify oomycete sRNAs that were expressed during infection and might be transferred into plants, we performed two types of sRNA-seq experiments. We on the one hand sequenced sRNAs isolated from total RNA extracts at 4 and 7 days post inoculation (dpi) together with mock-treated plants in order to find oomycete sRNAs expressed during infection. On the other hand, we sequenced sRNAs from *At*AGO1 immunopurification (*At*AGO1-IP) at 3 dpi to identify invasive oomycete sRNAs. We chose *At*AGO1 for the immunopurification sequencing, given that *At*AGO1 is constitutively expressed and forms the major RISC in *Arabidopsis* (Vaucheret, 2008), and because sRNAs of fungal pathogens were previously found to be associated with *At*AGO1 during infection (Wang et al., 2016; Weiberg et al., 2013).

We here describe the first sRNA transcriptome of *H. arabidopsidis* infecting *Arabidopsis*. An overview of total *Arabidopsis* and *Hyaloperonospora* sRNAs read number identified in all experiments is given in Tab.S1. Reads of total *Hpa*sRNAs were clustered in two major peaks of 21 nucleotides (nt) and 25 nt (Fig.1a), resembling sRNA size profiles previously reported for other plant pathogenic *Phytophthora* species (Fahlgren et al., 2013) suggesting that at least two categories of sRNAs occur in oomycetes. The identified *Hpa*sRNAs mapped to distinct regions of the *H. arabidopsidis* reference genome including ribosomal RNA (rRNA), transfer RNA (tRNA), small nuclear/nucleolar RNA, protein-coding genes (mRNA) and non-annotated regions (Fig.S1a). After filtering out rRNA, tRNA and snRNA reads, *Hpa*sRNAs mapping to protein-coding genes and non-annotated regions still displayed 21 nt as well as 25 nt size enrichment (Fig.S1b) and 5’ terminal uracil enrichment (Fig.1b).

**Figure 1:**
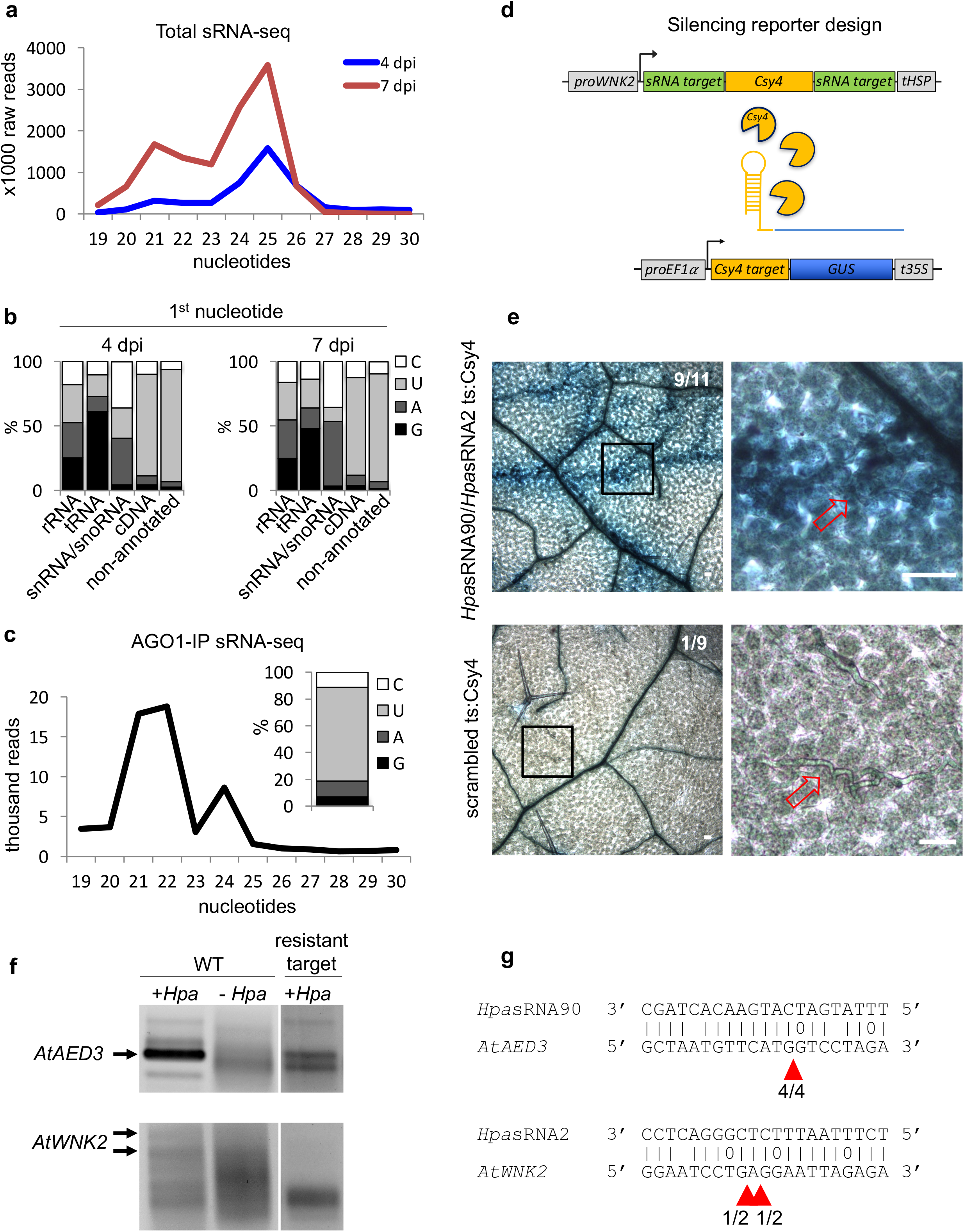
*Hpa*sRNAs invade the plant AGO1 and induce host target silencing in infected plant cells. a) Size profile of *Hpa*sRNAs revealed two size peaks at 21 nt and 25 nt at 4 and 7 dpi. b) The frequency of the first nucleotide at 5’ positions of *Hpa*sRNAs mapping to cDNAs or non-annotated regions revealed bias towards uracil. c) Size distribution and first nucleotide analysis of AtAGO1-associated *Hpa*sRNAs showed size preference at 21 nt with 5’ terminal uracil. d) A novel Csy4/GUS reporter construct was assembled to detect *Hpa*sRNA-directed gene silencing, reporting GUS activity if *Hpa*sRNAs were active to suppress Csy4 expression by sequence-specificity. e) GUS staining of infected leaves at two magnifications revealed sequence-specific reporter silencing at 4 dpi. Csy4 with *Hpa*sRNA2 and *Hpa*sRNA90 target sequence (ts) is depicted on the top and with scrambled ts on the bottom. Red arrows indicate *Hyaloperonospora* hyphae in the higher magnification images. Scale bars indicate 50 μm. f) 5’ RACE-PCR revealed amplification products at the expected size of target cleavage of *Hpa*sRNAs in infected WT plants, as indicated by arrows. Two PCR bands were detected for *AtWNK2* cleavage. No cleavage products corresponding to *Hpa*sRNAs were visible in non-infected WT plants for *AtAED3* and *AtWNK2* and for the resistant target version of *AtWNK2*. g) Sequencing of 5’ RACE-PCR products from infected WT plants revealed plant mRNA cleavage products at the *Hpa*sRNA target sequences, as indicated by red arrowheads.

We also identified *Hpa*sRNA reads in the *At*AGO1-IP sRNA-seq data providing first evidence that *Hpa*sRNAs translocated into plant cells and invaded the plant RISC. AtAGO1-associated *Hpa*sRNA reads highlighted 21 nt size enrichment with 5’ terminal uracil preference (Fig.1c), resembling typical profile of *Arabidopsis* AGO1-bound sRNAs (Fig. S1c) (Mi et al., 2008). This suggests that *Hpa*sRNAs were loaded into *At*AGO1. The ratio of *At*AGO1-bound *Hpa*sRNAs to total *Hpa*sRNAs was 1:78, whereas the ratio of the 21 nt *Hpa*sRNA fractions was 1:18 supporting that 21 nt *Hpa*sRNAs were preferably transferred into plant *At*AGO1. We suspected that such 21 nt *Hpa*sRNAs might have the potential to silence plant genes. Indeed, we identified 133 unique *Hpa*sRNA reads that were present in all infected total sRNA and AtAGO1-IP sRNA datasets with read counts > 5 reads per million in at least one dataset, among which 34 were predicted to target at least one *A. thaliana* cDNA with stringent cut-off criteria (Tab.S2).

### *Hpa*sRNAs induce host target silencing in infected plant cells

To examine if *At*AGO1-bound *Hpa*sRNAs could induce gene silencing in plants, we generated a novel *in situ* silencing reporter construct for *Hpa*sRNA-induced gene silencing. This reporter is based on the CRISPR endonuclease Csy4 that specifically binds and cleaves a short sequence motif (Haurwitz et al., 2010). This motif was fused to a *GUS* gene to mark it for degradation (Fig.1d). We cloned native plant target sequences of *Hpa*sRNAs as flanking tags next to the Csy4 to turn it into a target of *Hpa*sRNAs and to examine their potential of silencing the predicted plant targets. The GUS reporter was expected to become active if *Hpa*sRNAs would silence Csy4. For this examination, we chose as invading sRNA candidates *Hpa*sRNA2 and *Hpa*sRNA90 that were predicted to target *Arabidopsis AtWNK2* and *AtAED3* (Tab.S2). We chose these *Hpa*sRNA targets, because *AtWNK2* and *AtAED3* mRNAs were previously reported to accumulate less upon virulent *H. arabidopsidis* infection when compared to infection with an avirulent *H. arabidopsidis* strain by RNA-seq (Asai et al., 2014) supporting a potential virulence-triggered gene suppression. Both, *Hpa*sRNA2 and *Hpa*sRNA90 were confirmed to accumulate in infected plants at 4 and 7 dpi (Fig.S2). As promoter for Csy4, we cloned a 2 kb-DNA fragment upstream of the start codon of one of the two target genes (here *proAtWNK2*) to simulate native target mRNA levels. To exclude *Hpa*sRNA2/*Hpa*sRNA90-unspecific suppression of Csy4 or ckRNAi-independent effects that would activate GUS, we cloned scrambled target sequences of *Hpa*sRNA2 and *Hpa*sRNA90 as well as the target sequence of AtmicroRNA164 from its endogenous target *AtCUC2*, as negative controls. We tested at least three individual T1 lines per reporter construct. Csy4 blocked GUS activity in non-colonized cells (Fig.1e) proving functionality of the reporter repression. Upon infection, plants expressing Csy4 transcripts fused to *Hpa*sRNA2 and *Hpa*sRNA90 target sequences highlighted GUS activation along the *H. arabidopsidis* hyphal infection front (Fig.1e). This experiment provides, to our knowledge, the first *in situ* visualization of a pathogen’s sRNA translocation and function in infected host cells to trigger ckRNAi. GUS activation appeared only around the hyphae indicating that ckRNAi did not spread further into distal regions away from the primary infection. In contrast, Csy4 linked to scrambled *Hpa*sRNA2/*Hpa*sRNA90 target sequences or *At*miRNA164 target sequence did not typically express GUS activation around the infecting hyphae, excluding any target sequence-unspecific regulation of Csy4 or GUS, as well as pathogen-triggered regulation of the *AtWNK2* promoter (Fig.1e, Fig.S3).

As we revealed *Hpa*sRNA invasion into the plant *At*AGO1-RISC during infection and found infection-site specific target silencing triggered by *Hpa*sRNA2 or *Hpa*sRNA90, we attempted to clarify if *Hpa*sRNA2 and *Hpa*sRNA90 mediate gene silencing of its predicted plant targets for virulence. To test this, we performed quantitative reverse transcriptase (qRT)-PCR of AtWNK2 and AtAED3 on whole seedling leaves of wild type (WT) plants upon *H. arabidopsidis* infection and included the *atago1-27* mutant allele as a control, where target suppression should fail. Indeed, *AtAED3* was significantly down-regulated upon *H. arabidopsidis* inoculation at 7 dpi and *AtWNK2* expression indicated moderate suppression at 4 dpi (Fig.S4a) in WT, when compared to mock-treated samples. Because the down-regulation effects were moderate, we validated the results by a second independent *H. arabidopsidis* inoculation experiment (Fig.S4b). In support of *Hpa*sRNA-induced target silencing through *At*AGO1, suppression of *AtWNK2* and *AtAED3*, as observed in WT was abolished in the *atago1-27* background (Fig.S4). However, *AtAED3* expression data also indicated slight down-regulation upon mock treatment compared to before infection, as well as higher transcript levels in *atago1-27* before infection when compared to WT.

As *Arabidopsis* target transcripts were found to be down-regulated upon *H. arabidopsidis* infection, we examined, if *Hpa*sRNAs guided slicing of *AtWNK2* and *AtAED3* via the host *At*AGO1-RISC. *At*AGO1 possesses RNA cleavage activity on microRNA-guided target genes precisely at position 10/11 of the 5’end microRNA/mRNA duplex (Mallory and Bouché, 2008). We performed 5’ random amplification of cDNA-ends (RACE)-PCR analysis to reveal 5’ ends of target transcript fragments. We expected cleavage in *Hyaloperonospora-infected* WT plants but no cleavage products when using non-infected *Arabidopsis* or infected *Arabidopsis* expressing an *Hpa*sRNA-resistant version of *AtAED3r* or *AtWNK2r* (Fig.S5a). Indeed, using *Arabidopsis* WT resulted in PCR products of the expected cleavage size for both target genes (Fig.1f). While a clear band of the expected size was detected for *AtAED3*, two PCR bands were detected for *AtWNK2* at the expected size region in infected WT plant samples (Fig.1f). *AtWNK2* is predicted to have up to six splicing variants that would render RACE-PCR analysis being challenging and might explain for the two PCR bands. We found evidence for target mRNA cleavage at the predicted *Hpa*sRNA target sequence for both target genes by sequencing the isolated PCR products (Fig.1g). Cleavage was obtained slightly shifted from the predicted *At*AGO1 slicing position, namely at positions 8/9 opposite to 5’ *Hpa*sRNA90 and 10/11 and 11/12 to 5’ *Hpa*sRNA2. Indeed, we found alternative *Hpa*sRNA90 and *Hpa*sRNA2 species in our sRNA sequence libraries, which could explain the detected cleavage products as *At*AGO1-mediated. On the contrary, no PCR products of the expected size were obtained in uninfected plants and mutants with resistant target version of *At*WNK2 and in uninfected plants for *At*AED3, whereas AtAED3 resistant target version showed two bands slightly lower of the expected size (Fig.1g). Sequencing of cloned “nearby-expected size” PCR products in the *At*AED3 target resistant version revealed mRNA ends exclusively outside the predicted *Hpa*sRNA target sequence (Fig.S5b).

### *Arabidopsis atago1* exhibited enhanced disease resistance against downy mildew

Over hundred *Hpa*sRNAs invaded the plant AGO1/RISC during infection, with 34 *Hpa*sRNAs being predicted to silence 49 plant targets including stress-related genes (Tab.S2). *Hpa*sRNAs induced target host gene silencing at the infection site (Fig.1e). Based on these observations, we hypothesized that *At*AGO1 was crucial for *Hpa*sRNAs to suppress resistance. To test this hypothesis, we compared the disease outcome of *atago1-27* with WT plants by staining infected leaves with Trypan Blue. The hypomorphic *atago1-27* mutant represents relatively mild phenotypes compared to other *atago1* mutant alleles (Morel et al., 2002), enabling to perform infection assays. The *atago1-27* mutant exhibited a remarkable phenotype revealing dark stained host cells around hyphae, what we interpreted as trailing necrosis of plant cells (Fig.2a), a phenotype frequently observed in sub-compatible interactions (Coates and Beynon, 2010). This phenotype co-occurred with enhanced disease resistance, because *H. arabidopsidis* DNA content was strongly reduced (Fig.2b) and number of *H. arabidopsidis* conidiospores was significantly lower in *atago1-27*(Fig.2c). Pathogen DNA content was also reduced in *atago1-27* cotyledons (Fig.S6a) but without observing trailing necrosis (Fig.S6b), as previously described in sub-compatible combinations of *H. arabidopsidis* pathotypes and *A. thaliana* ecotypes (McDowell et al., 2005). The disease phenotype was indeed linked to *atago1* mutations and not to any secondary background mutation, as independent mutant alleles *atago1-45* and *atago1-46* also displayed, albeit to a smaller extent, trailing necrosis after inoculation with *H. arabidopsidis* (Fig.S6c). On the contrary, *atago2-1* and *atago4-2* did neither exhibit trailing necrosis nor reduced oomycete biomass (Fig.S6d,e). *Hpa*sRNA2 and *Hpa*sRNA90 were confirmed to load into *At*AGO1 but not into *At*AGO2 by AGO-IP coupled to stem-loop RT-PCR (Fig.S7), which is consistent with the observed reduced disease level in *atago1* mutants but not in *atago2-1*, suggesting that invasive *Hpa*sRNAs may act mainly through *At*AGO1 to support virulence.

**Figure 2:**
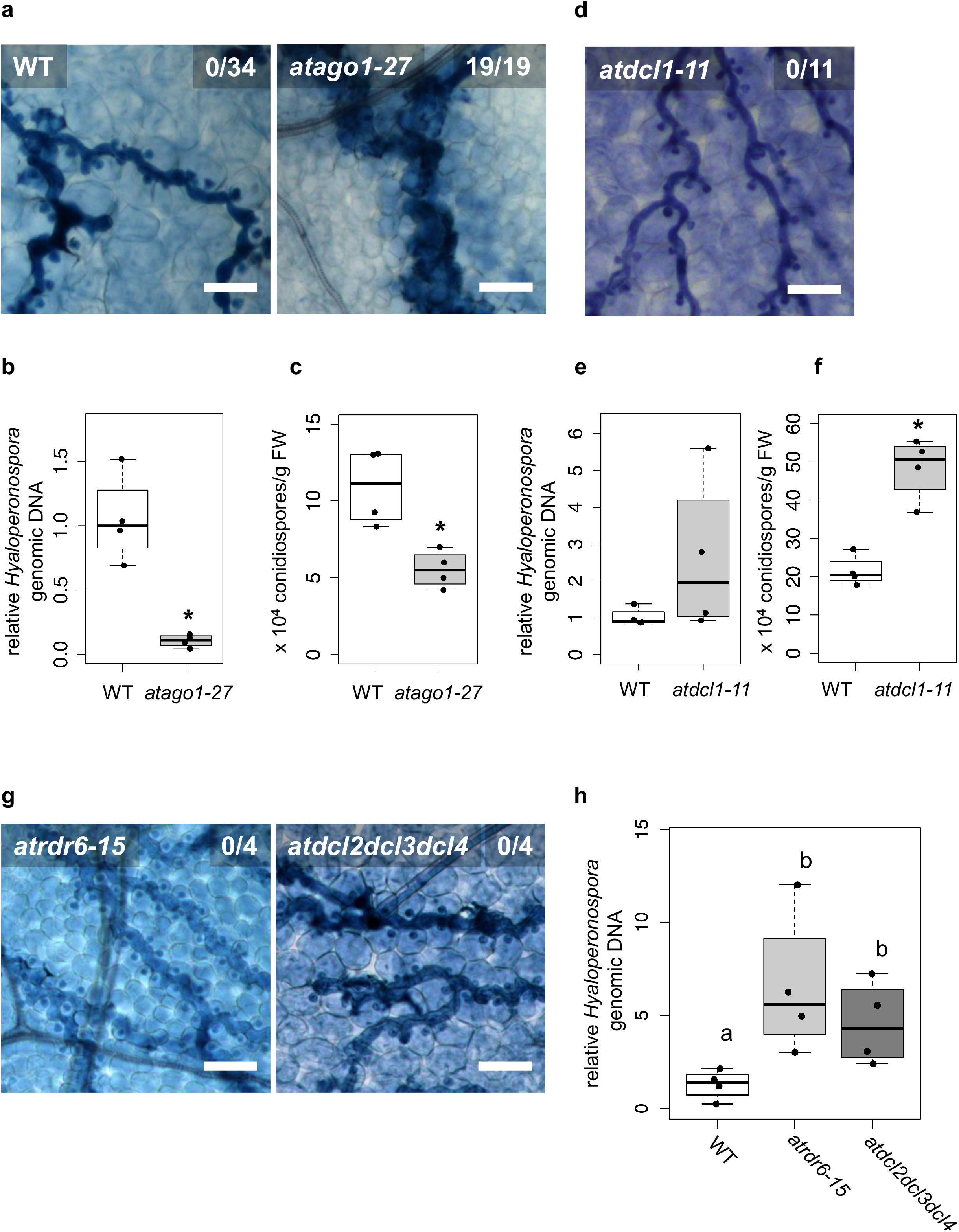
*Arabidopsis atago1* exhibited enhanced disease resistance against *H. arabidopsidis*. a) Trypan Blue-stained microscopy images showed trailing necrosis around hyphae in *atago1-27*, but no necrosis on WT seedling leaves at 7 dpi. b) *H. arabidopsidis* genomic DNA was quantified in *atago1-27* and WT plants by qPCR at 4 dpi relative to plant genomic DNA represented by n ≥ four biological replicates. Asterisk indicates statistically significant difference by one tailed Student’s test with p ≤ 0.05. c) Numbers of conidiospores per gram leaf fresh weight (FW) in *atago1-27* and WT plants at 7 dpi are represented by four biological replicates. The asterisk indicates significant difference by one tailed Student’s t-test with p ≤ 0.05. d) Trypan Blue-stained microscopy images of *atdcl1-11* did not show any trailing necrosis at 7 dpi. e) *H. arabidopsidis* genomic DNA in *atdcl1-11* and WT plants at 4 dpi were in tendency enhanced with n four biological replicates. f) Number of conidiospores per gram leaf fresh weight (FW) in *atdcl1-11* at 7 dpi was significantly elevated compared to WT plants. g) Trypan Blue-stained microscopy images of *atrdr6-15* and *atdcl2dcl3dcl4* showed no necrosis after inoculation with *H. arabidopsidis* at 7 dpi. h) *H. arabidopsidis* genomic DNA content in leaves was elevated in *atrdr6-15* and *atdcl2dcl3dcl4* compared to WT at 4 dpi with n ≥ four biological replicates. Letters indicate groups of statistically significant difference by ANOVA followed by TukeyHSD with p ≤ 0.05. Scale bars in all microscopy images indicate 50 μm and numbers represent observed leaves with necrosis per total inspected leaves.

The above results could have been also caused by debilitated plant endogenous sRNAs as *atago1* as well as other microRNA pathway mutants, such as *atdicer-like(dcl)1, athua enhancer(hen)1 athasty(hst)* or *atserrate(se)*, express developmental defects (Li and Zhang, 2016), which could have caused enhanced disease resistance against *H. arabidopsidis* (Fig.2a-c). To rule out this possibility, we inoculated *atdcl1-11* with *H. arabidopsidis*. We did not detect any trailing necrosis, reduced pathogen biomass, but even a significantly increased number of conidiospores in *atdcl1-11* (Fig.2d-f) indicating a positive role of *A. thaliana* microRNAs in immune response against *H. arabidopsidis*. These observations proved that necrotic trailing and reduced pathogen susceptibility found in *atago1* was not due to the loss of functional endogenous plant microRNA pathway. In support, *atse-2, athen1-5* and *athst-6* did also not show necrotic trailing upon infection (Fig.S8a,b).

Since *atago1* expressed trailing necrosis and reduced susceptibility to *H. arabidopsidis*, we wanted to rule out that common immunity or activation of *Resistance* (R) genes caused enhanced resistance in *atago1-27* compared to WT. We profiled gene expression of the *A. thaliana* immunity marker *AtPathogenesis-Related (PR)1*. Induction of *AtPR1* was neither faster nor stronger at 6, 12 or 18 hours post inoculation in *atago1-27* compared to WT (Fig.S9a). Expression of *AtPR1* and another immunity marker *AtPlant-Defensin (PDF)1.2* were not higher compared to WT before or after infection in *atago1-27* at 1, 4 or 7 dpi (Fig.S9b,c).

Plant microRNAs/*At*AGO1 are known to initiate the production of secondary phased siRNAs (phasiRNAs), which negatively control the expression of *NLR* (*Nucleotide-binding domain Leucine-rich Repeat*) class *R* genes (Li et al., 2012). The lack of phasiRNAs could in theory result in enhanced expression of *NLR*s and lead to higher resistance against *H. arabidopsidis*. PhasiRNA production depends on the *At*RNA-dependent RNA polymerase (RDR)6 and *At*DCL2/*At*DCL3/*At*DCL4 (Komiya, 2017). To rule out *R* gene-based enhanced resistance due to lack of phasiRNAs, we inoculated *atrdr6-15* and *atdcl2dcl3dcl4* mutants with *H. arabidopsidis* Noco2. Both mutants did not exhibit either trailing necrosis (Fig.2g) or reduced pathogen biomass (Fig.2h) upon inoculation with *H. arabidopsidis*.

In order to investigate whether *atago1-27* is more resistant to another biotrophic pathogen, we performed infection assays with the powdery mildew fungus *Erysiphe cruciferarum*. We did neither observe plant cell necrosis, nor a reduction in pathogen biomass compared to WT (Fig.S10a,b). These data confirmed that the observed resistance in *atago1* against *Hyaloperonospora* is not based on generally enhanced immunity or on *R* gene-mediated resistance.

### Invasive *Hyaloperonospora* sRNAs are crucial for virulence

As *Arabidopsis atago1* mutants displayed reduced susceptibility towards *H. arabidopsidis* infection and since *Hpa*sRNAs were invading the host *At*AGO1-RISC to silence plant target genes, we investigated how important *Hpa*sRNAs were for virulence. To assess the importance of *Hpa*sRNAs for virulence, we cloned and expressed a triple short tandem target mimic (STTM) in *Arabidopsis* to collectively scavenge three *Hpa*sRNAs: *Hpa*sRNA2, *Hpa*sRNA30 and *Hpa*sRNA90 (Fig.3a, Tab.S2). We included *Hpa*sRNA30 in the STTM, because it was predicted to target a homolog of *AtWNK2*, namely *AtWNK5*, and so we assumed that *Hpa*sRNA30/*At*WNK5 might also be important for virulence. *Hpa*sRNA30 was detectable in infected plant leaves at 4 and 7 dpi by stem-loop RT-PCR, but not at 0 and 1 dpi supporting that this sRNA was produced by *H. arabidopsidis* but not by *Arabidopsis* (Fig.S2a). Remarkably, seven out of eleven individual STTM T1 transgenic lines resembled partially the trailing necrosis phenotype also found in *atago1* (Fig.3b). We isolated two stable STTM T2 lines (#4, #5). The STTM #4 line showed target de-repression of *AtAED3* at 7 dpi and *AtWNK2* at 4 dpi when compared to plants expressing an empty vector control upon *H. arabidopsidis* inoculation (Fig.S11a). These time points corresponded to target gene suppression upon *H. arabidopsidis* inoculation, as found by qRT-PCR analysis (Fig. S4). Both STTM T2 lines exhibited reduced pathogen biomass (Fig.S11b) and allowed significantly lower production of pathogen conidiospores (Fig.3c). These results demonstrated that *Hpa*sRNAs are important for virulence.

**Figure 3:**
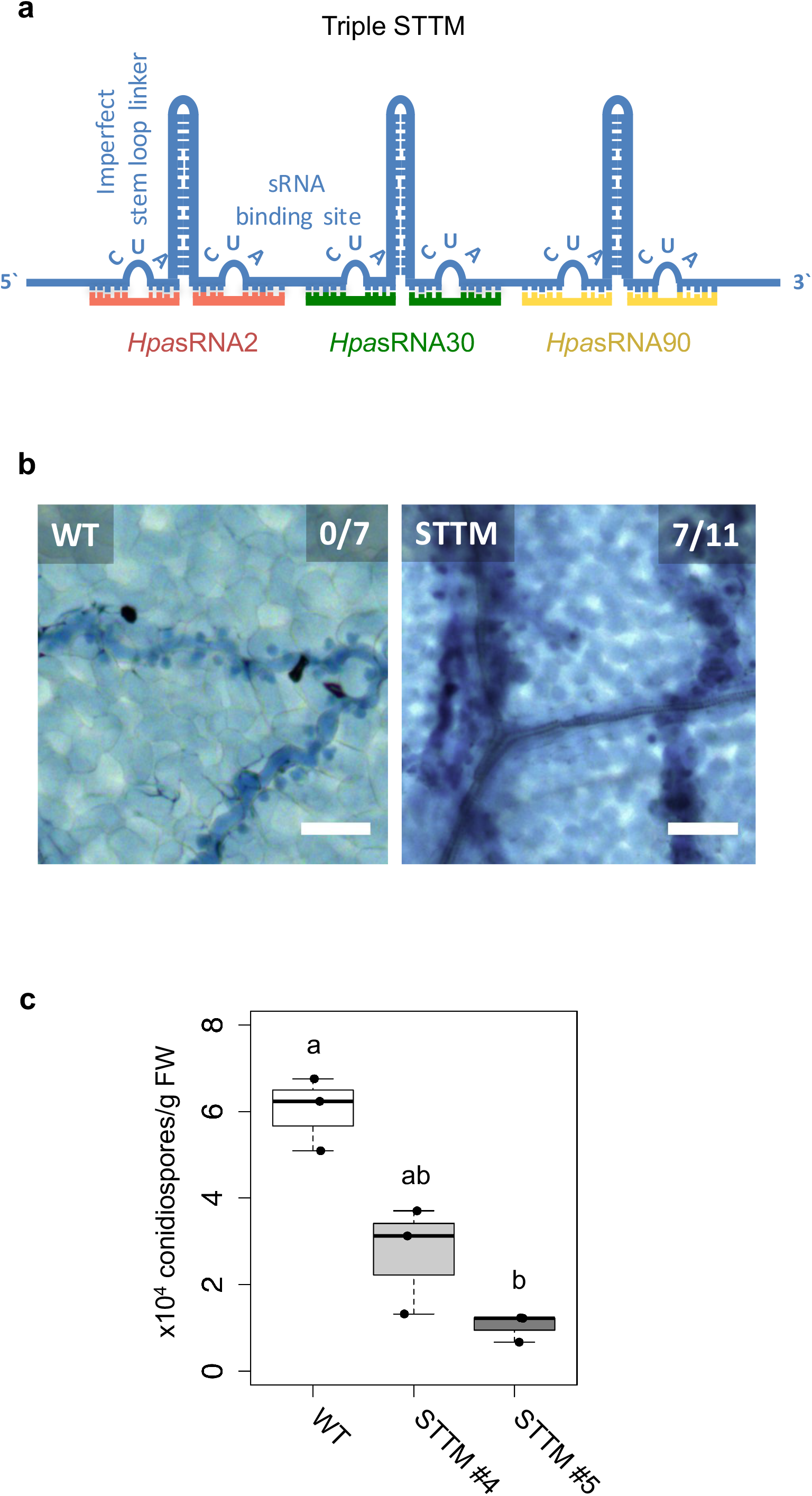
Invasive *Hpa*sRNAs are crucial for virulence. a) Triple STTM construct was cloned to target the three *Hpa*sRNAs *Hpa*sRNA2, *Hpa*sRNA30 and *Hpa*sRNA90 in *Arabidopsis*. b) *A. thaliana* T1 plants expressing the triple STTM to scavenge *Hpa*sRNA2, *Hpa*sRNA30 and *Hpa*sRNA90 exhibited trailing necrosis at 7 dpi. The scale bars indicate 50 μm and numbers represent observed leaves with necrosis per total inspected leaves. c) Number of conidiospores per gram FW was significantly reduced in two independent STTM-expressing *Arabidopsis* T2 lines (#4, #5) compared to WT. Letters indicate significant difference according to one site ANOVA including three biological replicates.

### *Arabidopsis* target genes of *Hyaloperonospora* sRNAs contribute to plant defence

To examine the contribution of individual target genes to plant defence, we isolated three T-DNA insertion lines, *atwnk2-2, atwnk2-3*, and *ataed3-1* (Fig.S12a), for inoculation with *H. arabidopsidis*. We located the T-DNA insertion in *atwnk2-3* from the last exon into the 3’ UTR based on sequencing of the T-DNA flanking site (Fig.S12a). Trypan Blue staining of infected leaves indicated WT-like infection structures in all T-DNA insertion lines. However, the number of haustoria at the conidiospore germination tube was significantly increased in *atwnk2-2* (Fig.S12b). The pathogen DNA content was slightly but not significantly enhanced in *atwnk2-2* and *ataed3-1* compared to WT, but not in *atwnk2-3* (Fig.4a). Significantly increased number of conidiospores (Fig.4b) and sporangiophores (Fig.4c) was found in all tested mutant lines upon *H. arabidopsidis* infection.

**Figure 4:**
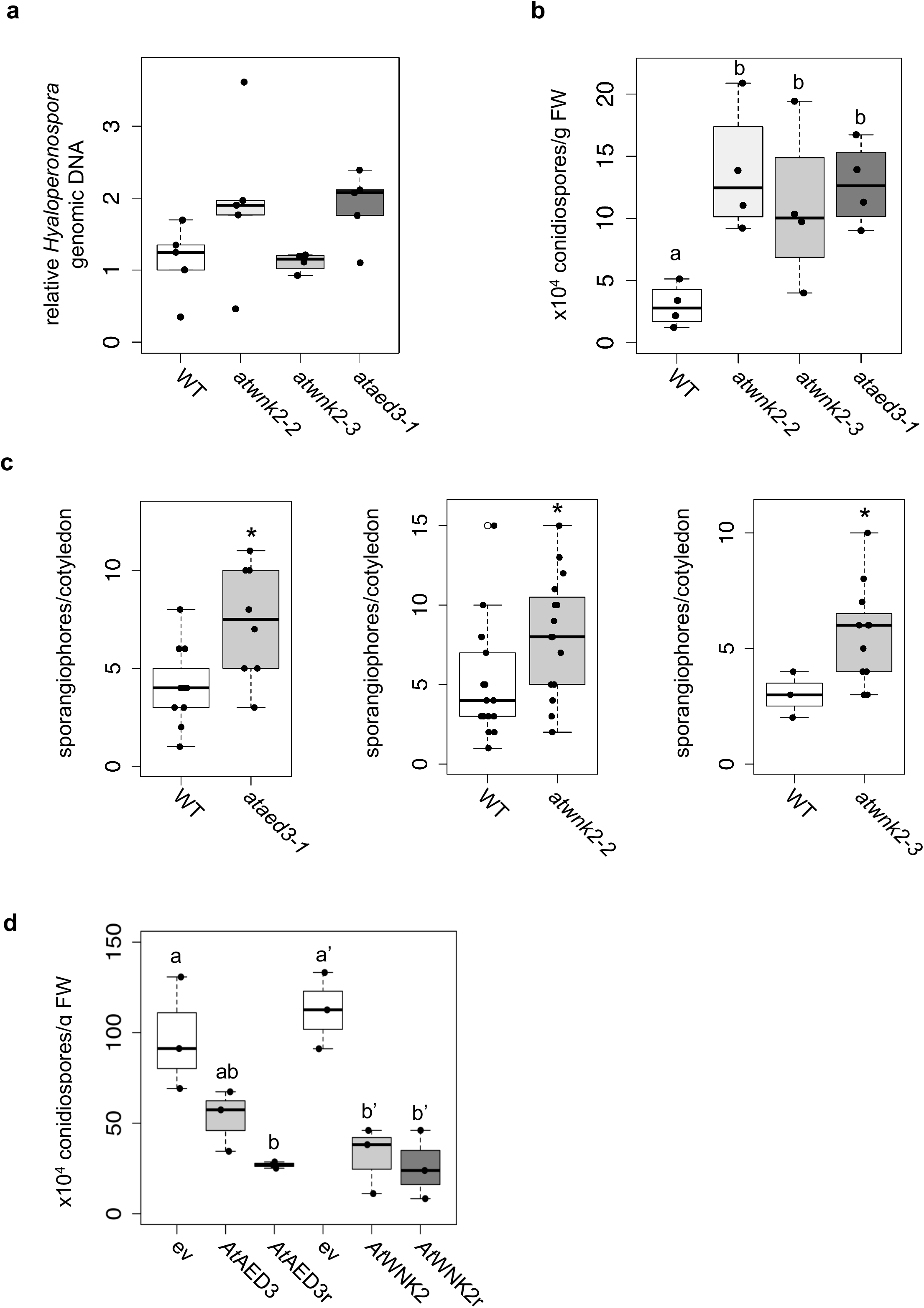
*Arabidopsis* target genes of *Hpa*sRNAs contribute to plant defence. a) *H. arabidopsidis* genomic DNA content in leaves was slightly but not significantly enhanced in *atwnk2-2* and *ataed3-1* compared to WT, but not in *atwnk2-3*, at 4 dpi with n ≥ four biological replicates. b) T-DNA insertion lines of *Hpa*sRNA target genes *ataed3-1, atwnk2-2*, and *atwnk2-3* showed significantly higher number of sporangiophores per cotyledon upon infection compared to WT at 5 dpi. Asterisks indicate significant difference by one tailed Student’s t-test with p ≤ 0.05. c) *ataed3-1, atwnk2-2*, and *atwnk2-3* showed significantly higher numbers of conidiospores per gram leaf FW upon infection compared to WT at 5 dpi. Letters indicate significant difference by one-site ANOVA test. d) Number of conidiospores was significantly reduced in complemented mutant lines using the native corresponding promoter (*proAtEWNK2, proAtAED3*) with native gene sequence, AtAED3 and AtWNK2, or with target site resistant version, AtAED3r and AtWNK2r compared to the knockout mutant background expressing an empty vector (ev), respectively. Letters indicate significant difference by one-site ANOVA test.

To continue the investigation on the contribution of individual target genes to plant defence, we expressed native promoter-driven (*proAtAED3, proAtWNK2*) targets *At*AED3 and *At*WNK2 or a target gene-resistant version, *At*AED3r and *At*WNK2r (Fig.S5a), in the respective mutant background *ataed3-1* and *atwnk2-2*. The integration of both *At*WNK2 and *At*WNK2r complemented the previously described early flowering phenotype of *atwnk2-2* (Wang et al., 2008) confirming that complementation of *At*WNK2 was successful (Fig. S13). We expected those plant lines to become more resistant against *H. arabidopsidis*. Indeed, both, the native gene version and the target site resistant version, exhibited reduced number of conidiospores compared to T-DNA mutant plants carrying an empty expression vector, respectively (Fig.4d). To further explore the role of target genes in plant immunity, we attempted to generate overexpression lines of resistant target gene versions and achieved an overexpressor of the *AtWNK2r* version (*AtWNK2*r-OE) (Fig.S5a) in the *atwnk2-2* background. *AtWNK2r*-OE plants showed ectopic cell death in distance from infection sites (Fig.S14a), as previously described for overexpression lines of other immunity factors, such as *At*BAK1 (Domínguez-Ferreras et al., 2015). Moreover, infection structures frequently displayed aberrant swelling-like structures and extensive branching of hyphae instead of the regular pyriform haustoria formed in *atwnk-2-2* (Fig.S14b), further supporting a role for *AtWNK2* in immune reaction.

## Discussion

Our study demonstrates the invasion, function, and significance of *Hyaloperonospora* sRNAs in virulence, the first natural ckRNAi case ever reported for an oomycete plant pathogen. By *Arabidopsis At*AGO1-IP coupled to sRNA-seq, we identified 34 *H. arabidopsidis* sRNAs that hijacked the host RNAi machinery to target multiple plant genes for silencing. These deep sequencing data offers first insights into the invasive *H. arabidopsidis* sRNA transcriptome during host infection.

By using a novel Csy4/GUS repressor reporter, oomycete sRNA-induced functional target gene silencing was demonstrated *in situ* and revealed effective silencing alongside the *Hyaloperonospora* hyphae. Compared to plant sRNAs, a relatively small proportion of *Hpa*sRNAs was counted in the AtAGO1 sRNA-seq experiment (1:2400), because most AtAGO1 molecules were purified from non-colonized tissue and *Hpa*sRNA-induced target silencing was only detected in *Arabidopsis* cells in close proximity to the pathogen hyphae. We suggest to not *pro forma* exclude sRNAs exhibiting low read number from ckRNAi studies, as other studies revealed strong phenotypic effects despite small RNA reads in the range of ten per million or lower (Jahan et al., 2015; Qutob et al., 2013). *At*AGO1 was a major RISC that was hijacked by *Hpa*sRNAs to success infection, because both blocking *Hpa*sRNAs by transgenic target mimics and dysfunctional *atago1* mutant alleles displayed a clear disease resistance phenotype. We investigated two examples of *Hpa*sRNA target genes, *AtWNK2* and *AtAED3*, and confirmed mRNAs were down-regulated upon infection and cleaved at the predicted *Hpa*sRNA target sequences. Target genes suppression was moderate, as expected, since RNA was purified from whole leaves with most cells being from non-infected leaf lamina. This was confirmed by the results of the Csy4/GUS repressor reporter, that demonstrated that target silencing occurred only in the *Arabidopsis* cells in close proximity to the *Hyaloperonospora* hyphae and haustoria (Fig.1e).

Plant invasive *Hpa*sRNAs are crucial for successful infection, because transgenic *Arabidopsis* that block function of three examples *Hpa*sRNA2, *Hpa*sRNA30 and *Hpa*sRNA90 via target mimics diminished virulence. As we identified 113 invasive *Hyaloperonospora* sRNAs with 49 predicted plant target genes, we suggest that many *Hpa*sRNAs collaboratively sabotage expressional activation of plant immune response, as previously shown for proteinaceous pathogen effectors (Cunnac et al., 2011). Interestingly, we found that the *Hpa*sRNA2 sequence was conserved among plant pathogenic oomycete species of the genera *Hyaloperonospora*, *Phytophthora* and *Pythium* (Fig.S15a). Target sequences within plant *WNK2* homologs were conserved as well, with the lowest number of base pair mismatches occurring in the highly-adapted *A. thaliana/H. arabidopsidis* interaction (Fig.S15b), reflecting the most co-adapted plant-oomycete interaction. It will be exciting to see how sRNAs of other oomycete pathogens contribute to host infection.

Regarding the role of identified *Hpa*sRNAs target genes in host defence, our data supported quantitative contributions of *AtAED3* and *AtWNK2* to plant immunity. *AtAED3* encodes a putative apoplastic aspartyl protease and has been suggested to be involved in systemic immunity (Breitenbach et al., 2014). *AtWNK2* contributes to flowering time regulation in *A. thaliana*, while other members of the plant WNK family have been linked to the abiotic stress response (Cao-Pham et al., 2018). What is the particular function of these target genes against *H. arabidopsidis* infection and whether these also play a role against other pathogens, needs to be investigated.

With this new data, we demonstrate that ckRNAi is a conserved virulence mechanism among distinct classes of pathogens, as fungi deliver sRNAs into host AGO as well to target host genes (Weiberg et al., 2013), and ckRNAi should be considered in virtually all host-pathogen interactions.

## Author contributions

Research concept and design A.W.; experimental design: F.D., and A.W.; experiments performed F.D, A.T., J.S.R., S. K., R.P.; bioinformatics analysis: A.W. and F.D.; contribution of the Csy4-based de-repression system: T.S. and A.T.; manuscript writing: A.W. and F.D.; manuscript reviewing and editing: F.D., R.H., and A.W.

## Acknowledgements

The authors thank Michaela Pagliara for excellent technical assistance, Dr. Martin Parniske for critical reading of the manuscript and inspiring scientific discussions and support, and Dr. Aline Banhara and Fang-Yu Hwu for introducing us into the *Hyaloperonospora/Arabidopsis* pathosystem and the Gene Center Munich for Illumina sequencing service. Seeds used in this study were provided by the Nottingham *Arabidopsis* Stock Centre (NASC) unless otherwise specified. We thank Dr. Hervé Vaucheret, Dr. James Carrington, Dr. Steven Jacobsen, and for kindly providing us seeds of the *atago1-27, atdcl2dcl3dcl4, atrdr6-15, atse-2* mutants and Dr. Tino Köster for the *atdcl1-11* mutant. We thank Dr. Michael Boshart for providing us αHA (12CA5) antibody. We thank Dr. David Chiasson and Martin Bircheneder for providing Golden Gate entry plasmids. This work was supported by the German Research Foundation (DFG; Grant-ID WE 5707/1-1). The funders had no role in study design, data collection and analysis, and decision to publish or in preparation of the manuscript.

## Supplemental figure legends

Figure S1: a) *Hpa*sRNAs mapped to distinct coding and to non-coding genomic regions. b) Relative read counts and size distribution of *Hpa*sRNAs mapped to different genomic regions at 4 and 7 dpi. c) Size distribution and first nucleotide analysis of AtAGO1-associated sRNAs of *A. thaliana*.

Figure S2: Stem-loop RT-PCR revealed *Hpa*sRNA2, *Hpa*sRNA30 and *Hpa*sRNA90 expression at 4 and 7 dpi in three biological replicates. Total RNA served as loading control.

Figure S3: Csy4 repressor reporter with *Hpa*sRNA2 and *Hpa*sRNA90 target sequence (ts) is depicted on the left and with AtmiRNA164 ts of the *AtCUC2* target gene on the right. GUS staining of infected leaves at two magnifications revealed sequence-specific reporter silencing at 4 dpi in *Hpa*sRNA2/*Hpa*sRNA90 ts construct but not in AtmiRNA164 ts. Red arrows indicate *Hyaloperonospora* hyphae in the higher magnification images. Scale bars indicate 50 μm.

Figure S4: Relative expression of *AtAED3* and *AtWNK2* was measured in mock-treated or *H. arabidopsidis-infected* WT and *atago1-27* seedlings before and at 4 and 7 dpi by qRT-PCR using *AtActin* as a reference in two independent infection experiments (a, b). Numbers below graph give change fold factors of *Hyaloperonospora-infected* versus mock-treated samples. The bar represents the average of n ≥ three biological replicates. Numbers give change fold factors of *H. arabidopsidis-infected* divided by mock-treated samples, and letters indicate groups of statistically significant difference within one time point by ANOVA followed by TukeyHSD with p ≤ 0.05.

Figure S5: a) Target sequence-resistant versions of *AtAED3* (*AtAED3r*) and *AtWNK2* (*AtWNK2r*) were cloned by introducing synonymous nucleotide exchanges indicated by red letters. b) RACE-PCR sequencing *AtAED3r* in *Hyaloperonospora-infected* plants, as indicted by red arrowheads, did not match with the predicted *Hpa*sRNA target sequence (green box).

Figure S6: a) *H. arabidopsidis* genomic DNA content in cotyledons was lower in *atago1-27* compared to WT, as quantified by qPCR relative to plant genomic DNA at 4 dpi with n ≥ three biological replicates. Asterisk indicates significant difference by one tailed Student’s t-test with p ≤ 0.05. b) Trypan Blue-stained microscopy images of *atago1-27* cotyledons did not show any necrosis at 7 dpi. c) Trypan Blue-stained microscopy images of *atago1-45* and *atago1-46* revealed trailing necrosis at 7 dpi with *H. arabidopsidis*. d) Trypan Blue-stained microscopy images presenting *H. arabidopsidis-infected atago2-1* and *atago4-2* seeding leaves at 7 dpi. e) *H. arabidopsidis* genomic DNA was quantified in WT versus *atago2-1* and *atago4-2* by qPCR at 4 dpi relative to plant genomic DNA represented by n ≥ four biological replicates. Letters indicate no statistical difference by ANOVA test.

Figure S7: Stem-loop RT-PCR of *Hpa*sRNAs from *At*AGO1 or *At*AGO2 co-IP with mock-treated or *H. arabidopsidis-inoculated* seedlings. *At*miRNA164 and *At*miRNA393* were used as positive AtAGO co-IP controls. Pull-down of *At*AGO1 was achieved with WT plants using an *At*AGO1 native antibody, and *At*AGO2 with HA-epitope tagged *At*AGO2-expressing *A. thaliana* Col-0 using anti-HA antibody with the lower panel showing Western blot analysis.

Figure S8: Trypan Blue-stained microscopy images presenting the AtmiRNA biogenesis mutants *athst-6, athen1-5* and *atse-2* did not show any trailing necrosis at 7 dpi. Scale bars in microscopy images indicate 50 μm and numbers represent observed leaves with necrosis per total inspected leaves.

Figure S9: a) Expression analysis of *AtPR1* by RT-PCR in WT and *atago1-27* did not show obvious differences at 6 and 12 hours post inoculation with *H. arabidopsidis. AtActin* was used as reference gene with four biological replicates. b and c) Relative expression of *AtPR1* and *AtPDF1.2* determined by qRT-PCR using *AtActin* as reference. The bar represents the average of n ≥ three biological replicates, each comprising two technical replicates.

Figure S10: a) Trypan Blue-stained microscopy images of WT or *atago1-27* leaves infected with *Erysiphe cruciferarum* did not show necrosis at 8 dpi. Scale bars in microscopy images indicate 50 μm and numbers represent observed leaves with necrosis per total inspected leaves. b) *Erysiphe* genomic DNA content in WT and *atago1-27* was not significantly different at 4 dpi by qPCR relative to plant genomic DNA with n ≥ three biological replicates as determined by one tailed Student’s t-test.

Figure S11: a) Relative expression of *AtAED3* at 7 dpi and *AtWNK2* at 4 dpi was determined for STTM or empty vector (ev) expressing plants upon *H. arabidopsidis* inoculation at 7 and 4 dpi respectively by qRT-PCR. One biological replicate represents three leaves, the bar represents the average of n ≥ three biological replicates. b) *H. arabidopsidis* genomic DNA content in leaves was increased in STTM #4 and STTM #5 plants compared to empty vector (ev) expressing WT plants at 4 dpi with n ≥ three biological replicates.

Figure S12: a) T-DNA insertion of *Hpa*sRNA target gene mutant lines *atwnk2-2, atwnk2-3* and *ataed3-1* were verified by genomic DNA PCR. b) Trypan Blue-stained microscopy images revealed a higher number of haustoria in the first 200 μm of hyphae (indicated by white bar alongside the hyphae) from the spore germination site in *atwnk2-2* compared to WT with n ≥ eight leaves. Asterisk indicates significant difference by one tailed Student’s t-test with p ≤ 0.05. Similar results were obtained in two independent experiments.

Figure S13: Transgenic *A. thaliana atwnk2-2* was complemented with *proWNK2:WNK2* or *proWNK2:WNK2r* that resulted in a WT-like flowering time point, while empty vector (ev) exhibited early flowing phenotype, as reported for *atwnk2-2* (Wang et al., 2008).

Figure S14: a) *A. thaliana* plant expressing *AtWNK2r* in the *atwnk2-2* background revealed local necrosis without pathogen infection and b) aberrant hyphae and haustoria swellings. Scale bars in microscopy images indicate 50 μm and the numbers represent observed leaves with necrosis or swellings respectively per total inspected leaves.

Figure S15: a) Oomycete *SRNA2* genomic loci are conserved among different plant pathogenic oomycete species of the genera *Hyaloperonospora, Phytophthora*, and *Pythium* (Hpa = *Hyaloperonospora arabidopsidis*, Pcap = *Phytophthora capsici*, Pso = *Phytophthora sojae*, Pan = *Pythium aphanidermatum*, Pinf = *Phytophthora infestans*, Ppa = *Phytophthora parasitica*). Blue box at the consensus sequence indicates the region of sRNA transcription as identified by deep sequencing analysis and red box marks the consensus of the mature 21 nt *Hpa*sRNA2 region. b) Target prediction alignment of *sRNA2* homologs from different oomycete species with the target sequences of homolog *WNK2*s from respective host plant species (At = *Arabidopsis thaliana*, Gm = *Glycine max*, St = *Solanum tuberosum*, Nt = *Nicotiana tabacum)*. Double point indicates a base pair match, single point is a wobble base pair, and no-point represents a mismatch.

## Supplemental Tables

Table S1: sRNA read numbers

Table S2: Predicted *A. thaliana* target genes of *Hpa*sRNAs

Table S3: List of oligonucleotides used in this study

## Material & Methods

### Plant material and inoculation with *H. arabidopsidis*

*Arabidopsis thaliana* (L.) seedlings were grown on soil under long day condition (16 h light/8 h dark, 22 °C, 60 % relative humidity). The *atago1-27, atago1-45, atago1-46, atago2-1, atago4-2, athst-6, athen1-5, atse-2, atdcl1-11, atdcl2dcl3dcl4, atrdr6-15* (all in the Col-0 background) were described previously (Agorio and Vera, 2007; Allen et al., 2004; Bollman et al., 2003; Deleris et al., 2006; Grigg et al., 2005; Morel et al., 2002; Smith et al., 2009; Takeda et al., 2008; Vazquez et al., 2004; Zhang et al., 2008). The *atwnk2-2* (SALK_121042, (Wang et al., 2008)), *atwnk2-3* (SALK_206118) and *ataed3-1* (SAIL_722 G02C1) lines were verified for the T-DNA insertion by PCR on genomic DNA. *Hyaloperonospora arabidopsidis* (Gäum.) isolate Noco2 was maintained on Col-0 plants. Plant inoculation was performed using 2-2.5 × 10^4^ spores/ml and incubated in a growth chamber under long day condition at 18 °C as described previously (Ried et al., 2019). For *atwnk2-2, atwnk2-3, and ataed3-1* pathogen assays inoculum strength was reduced to 1 × 10^4^ spores/ml.

### Powdery mildew inoculation

*Erysiphe cruciferarum* was maintained on highly susceptible Col-0 *phytoalexin deficient (pad)4* mutants in a growth chamber at 22 °C, a 10-hour photoperiod with 150 μmol m^−2^s^−1^, and 60 % relative humidity. For pathogen assays 6-weeks old *Arabidopsis* plants were inoculated with *E. cruciferarum* in a density of 3–5 spores mm^−2^ and replaced under the same conditions.

### Trypan Blue staining

Infected leaves were stained with Trypan Blue as described previously (Koch and Slusarenko, 1990). Microscopic images were taken with a DFC450 CCD-Camera (Leica) on a CTR 6000 microscope (Leica Microsystems).

### GUS staining

Infected leaves were vacuum-infiltrated with GUS staining solution (0.625 mg ml^−1^ X-Gluc, 100 mM phosphate buffer pH 7.0, 5 mM EDTA pH 7.0, 0.5 mM K3[Fe(CN)6], 0.5 mM K_4_[Fe(CN)_6_], 0.1% Triton X-100) and incubated over night at 37°C. Leaves were de-stained with 70% ethanol overnight and microscopic images were taken with the same set up as Trypan Blue stained samples.

### Pathogen quantification

*Hyaloperonospora* spores were harvested at 7 dpi into 2 ml of water. Spore concentration was determined using a haemocytometer (Neubauer improved, Marienfeld). Number of sporangiophores were counted on detached cotyledons using a binocular. Genomic DNA was isolated using the CTAB method followed by Chloroform extraction and isopropanol precipitation (Chen and Ronald, 1999). Four leaves were pooled for one biological replicate and isolated DNA was diluted to a concentration of 5 ng μl^−1^. *H. arabidopsidis* and *A. thaliana* genomic DNA were quantified by qPCR on a qPCR cycler (CFX96, Bio-Rad) using SYBR Green (Invitrogen, Thermo Fischer Scientific) and GoTaq G2 Polymerase (Promega) using species-specific primers (Tab.S3). Relative DNA content was calculated using the 2^−ΔΔCt^ method (Livak and Schmittgen, 2001).

### *A. thaliana* gene expression analysis

Total RNA was isolated using a CTAB-based method (Bemm et al., 2016). Genomic DNA was removed using DNase I (Sigma-Aldrich) treatment and cDNA synthesis was performed with 1 μg total RNA using SuperScriptIII RT (Thermo Fisher Scientific). Gene expression was measured by qPCR on a qPCR cycler (CFX96, Bio-Rad) using SYBR Green (Invitrogen, Thermo Fisher Scientific) and GoTaq G2 Polymerase (Promega). Differential expression was calculated using the 2^−ΔΔCt^ method (Livak and Schmittgen, 2001).

### Generation of transgene expression vectors

Plasmids for *Arabidopsis* transformation were constructed using the plant Golden Gate based toolkit (Binder et al., 2014). The coding sequences of *AtWNK2* and *AtAED3* were amplified by PCR from *Arabidopsis* cDNA, and silent mutations were introduced by PCR in the target sequence of *Hpa*sRNA2 and *Hpa*sRNA90, respectively. For overexpression, *AtWNK2r* was ligated into a binary expression vector with a C-terminal YFP tag under the control of the *LjUbq1* promoter. *AtWNK2r* and *ATAED3r* were also ligated into a binary expression vector with a C-terminal *YFP* tag under the control of their native promoters (~2 kb upstream of the translation start site). Promoter function was tested by fusion to 2x*GFP-NLS* and fluorescence microscopy of transiently transformed *Nicotiana benthamiana* leaves. STTM sequences were designed as described previously (Tang et al., 2012), and flanks with BsaI recognition sites were introduced. STTM sequences were synthesized as single stranded DNA oligonucleotides (Sigma Aldrich). The strands were end phosphorylated by T4 polynucleotide kinase (NEB), annealed, and cloned into an expression vector under the control of the *pro35S*, and the final vector with STTMs for *Hpa*sRNA2, *Hpa*sRNA30, and *Hpa*sRNA90 in a row after each other was assembled. The coding sequence of Csy4 was synthesized (MWG Eurofins) with codon optimization for expression in plants. Cloned Csy4 was flanked with new overhangs for integration in the Golden Gate toolkit by PCR. A fusion of the target sequences of *Hpa*sRNA2 and *Hpa*sRNA90, the target sequence of AtmiRNA164a, and the target sequence of Csy4 were synthesized as single strands (Sigma Aldrich). The strands were end phosphorylated by T4 polynucleotide kinase (NEB) and annealed. Csy4 was flanked with the respective target sequences and ligated into a vector under the control of the *AtWNK2* promoter by BsaI cut ligation. For the reporter, a Csy4 target sequence was inserted between the Kozak sequence and the start codon of the *GUS* gene and ligated into a vector under the control of the *AtEF1α* promoter. The final binary expression vector was assembled by combination of the Csy4 and the *GUS* vectors by BpiI cut ligation. All cloning primers are listed in Tab.S3.

### Generation of transgenic *Arabidopsis* plants

*Arabidopsis* plants of Col-0 (WT), *atwnk2-2*, and *ataed3-1* were transformed with the respective construct using the *Agrobacterium tumefaciens* strain AGL1 by the floral dip method (Clough and Bent, 1998). Transformed plants were selected on ½ MS + 1% sucrose agar plates containing 50 μg/ml kanamycin, and were subsequently transferred to soil. Experiments were carried out on T1 generation plants representing independent transformants, unless a transformation line number is indicated (e.g. STTM #4). These experiments were carried out using T2 plants.

### AGO Western blot analysis and sRNA co-immunopurification

*A. thaliana* AGO1 or HA-tagged *At*AGO2 were co-immunopurified (co-IPed) and isolated as described previously (Zhao et al., 2012). 1 μg α-AGO1 antibody (Agrisera)/g leaf tissue or 0.1 μg α-HA antibody (3F10, Roche or 12CA5)/g leaf tissue were used. For Western blot analysis 30 % of the co-IP fraction were used, and protein was detected using α-AGO1 antibody (Agrisera) in 1:4000 dilution or α-HA antibody (3F10, Roche or 12CA5) in 1:1000 dilution, respectively. This was followed by an incubation with adequate secondary antibody (α-rabbit IRdye800 (LI-COR, 1:3000 dilution), α-mouse IRdye800 (LI-COR, 1:15000 dilution), and α-rat IRdye800 (LI-COR, 1:15000 dilution), and protein detection was performed with the Odyssey imaging system (LI-COR). Recovery of the co-IPed sRNAs was achieved as previously described (Carbonell et al., 2012), and was directly used for RT-PCR analysis or sRNA library cloning.

### Stem loop RT PCR

sRNAs were detected by stem-loop RT-PCR from 1 μg of total RNA or 5 % of the AtAGO co-IPed RNA, as described previously (Varkonyi-Gasic et al., 2007).

### 5’ RACE-PCR

5’ RACE-PCR was performed on 1 μg of total isolated from non-inoculated or *Hyaloperonospora*-infected *Arabidopsis* leaves pooled from equal amounts isolated at 4 and 7 dpi, using the 5’/3’ RACE Kit, 2nd Generation (Roche Diagnostics). After the first round of PCR, a band of the expected size was cut out and a nested PCR was carried out on the eluted DNA. Bands were cut out and DNA was eluted using GeneJet Gel Extraction Kit (Thermo Scientific). PCR fragments were blunted using Klenow fragment (NEB) and cloned into pUC57 and sequenced.

### sRNA cloning, sequencing and target gene prediction

sRNAs were isolated for high throughput sequencing as previously described (Weiberg et al., 2013). SRNAs were cloned for Illumina sequencing using the Next® Small RNA Prep kit (NEB) and sequenced on an Illumina HiSeq1500 platform. The Illumina sequencing data were analysed using the GALAXY Biostar server. Raw data were de-multiplexed (Illumina Demultiplex, Galaxy Version 1.0.0) and adapter sequences were removed (Clip adaptor sequence, Galaxy Version 1.0.0). Sequence raw data are deposited at the NCBI SRA server (BioProject accession: PRJNA395139). Reads were then mapped to a master genome of *Hyaloperonospora arabidopsidis* comprising the isolates Emoy2 (BioProject PRJNA30969), Cala2 (BioProject PRJNA297499), Noks1 (BioProject PRJNA298674) using the BOWTIE algorithm (Galaxy Version 1.1.0) allowing zero mismatches (-v 0). Subsequently, reads were cleaned from *Arabidopsis thaliana* sequences (TAIR10 release) with maximal one mismatch. For normalization, ribosomal RNA (rRNA), transfer RNA (tRNA), small nuclear RNAs (snRNAs), and small nucleolar RNA (snoRNA) reads were filtered out using the SortMeRNA program (Galaxy Version 2.1b.1). The remaining reads were count and normalized on total *H. arabidopsidis* reads per million (RPM). The *Hpa*sRNAs were clustered if their 5’ end position or 3’ end position were within the range of three nucleotides referring to the genomic loci (Weiberg et al., 2013). Target gene prediction of sRNAs was performed with the TAPIR program using a maximal score of 4.5 and a free energy ratio of 0.7 as thresholds (Bonnet et al., 2010).

### DNA alignment

Search for homologous sequences of *Hpa*sRNA2 was performed by BLASTn search using the Ensembl Protists database (http://protists.ensembl.org). Homologs DNA sequences of 100 nucleotides up- and downstream of *SRNA2* homologs were aligned using the CLC Main Workbench package.

### Statistical analysis

All statistical tests were carried out using R studio (version 1.0.136, rstudio.com). ANOVA tests were done on log-transformed data.

