## supplementary figures for "Oomycete small RNAs invade the plant RNA-induced silencing complex for virulence"

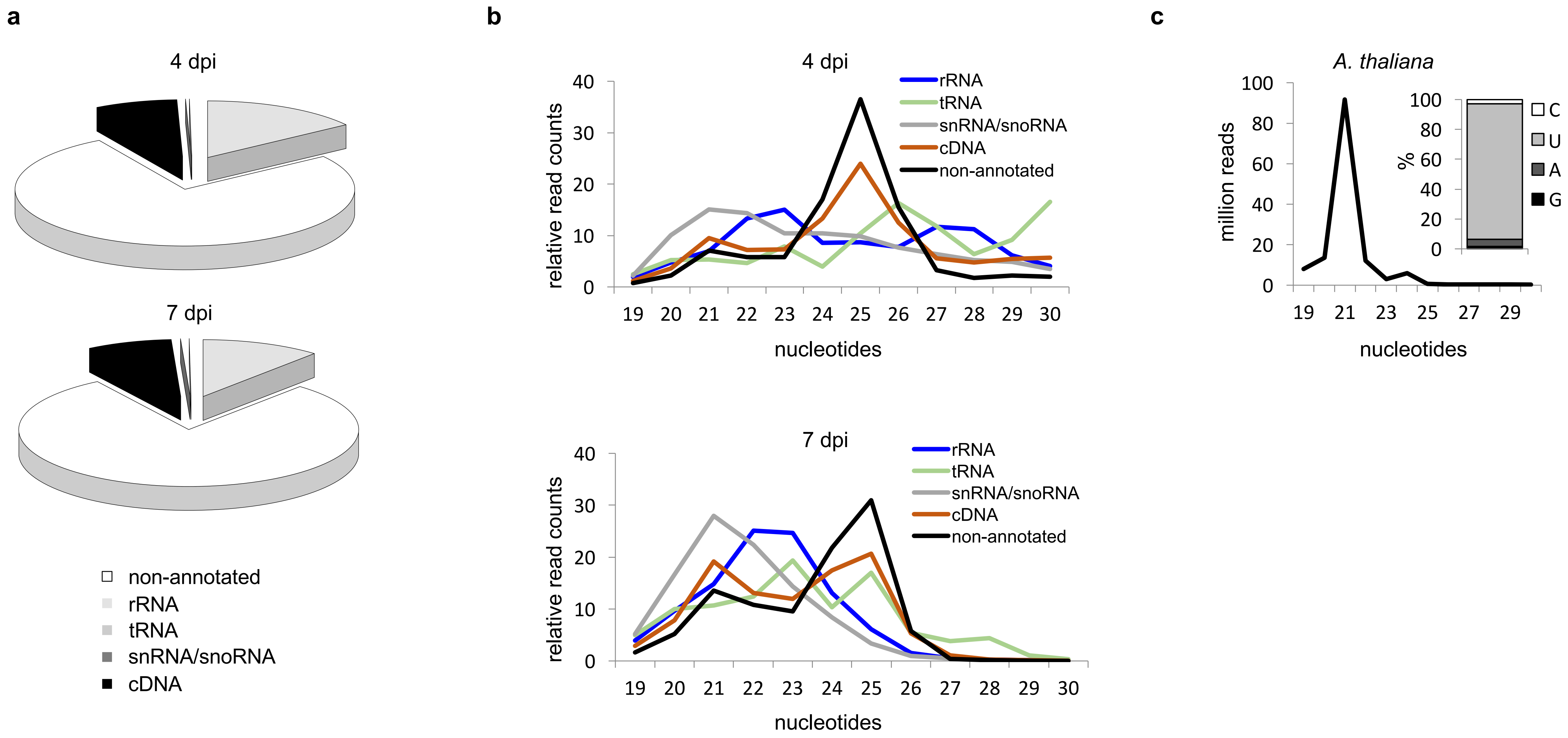
a
4 dpi
7 dpi
non-annotated
rRNA
tRNA
snRNA/snoRNA
cDNA
b
4 dpi
7 dpi
rRNA
tRNA
snRNA/snoRNA
cDNA
non-annotated
relative read counts
nucleotides
c
A. thaliana
million reads
%
C
U
A
G
nucleotides

**Figure S1**

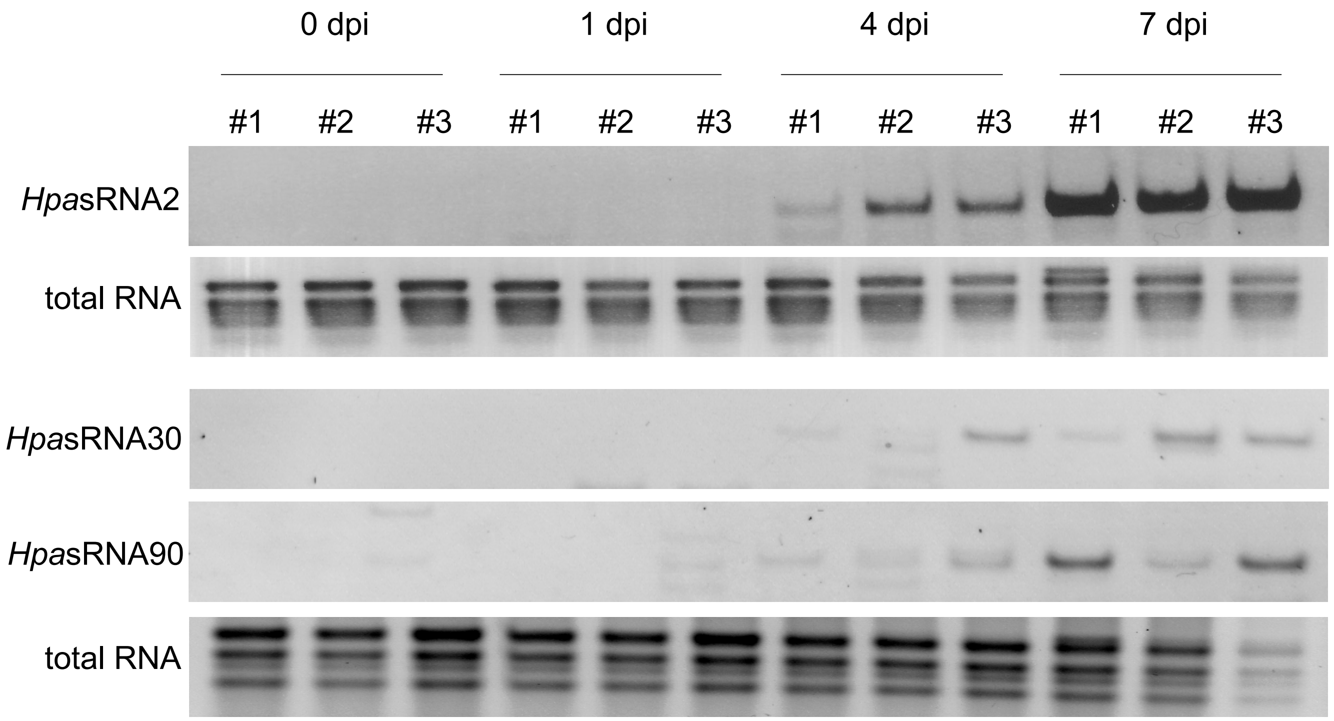
0 dpi
1 dpi
4 dpi
7 dpi
#1
#2
#3
*Hpa*sRNA2
total RNA
*Hpa*sRNA30
*Hpa*sRNA90
total RNA

**Figure S2**

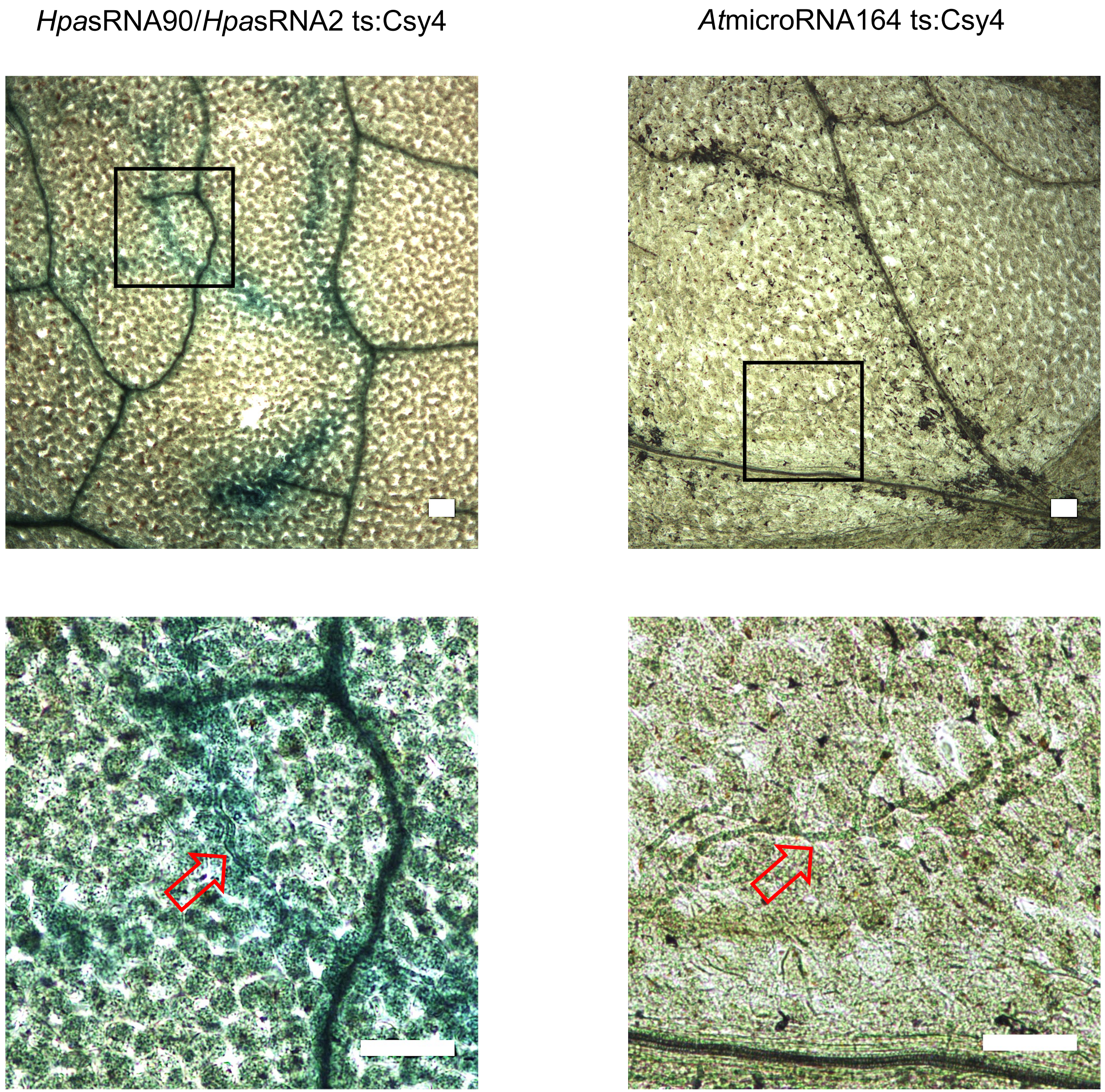


**Figure S3**

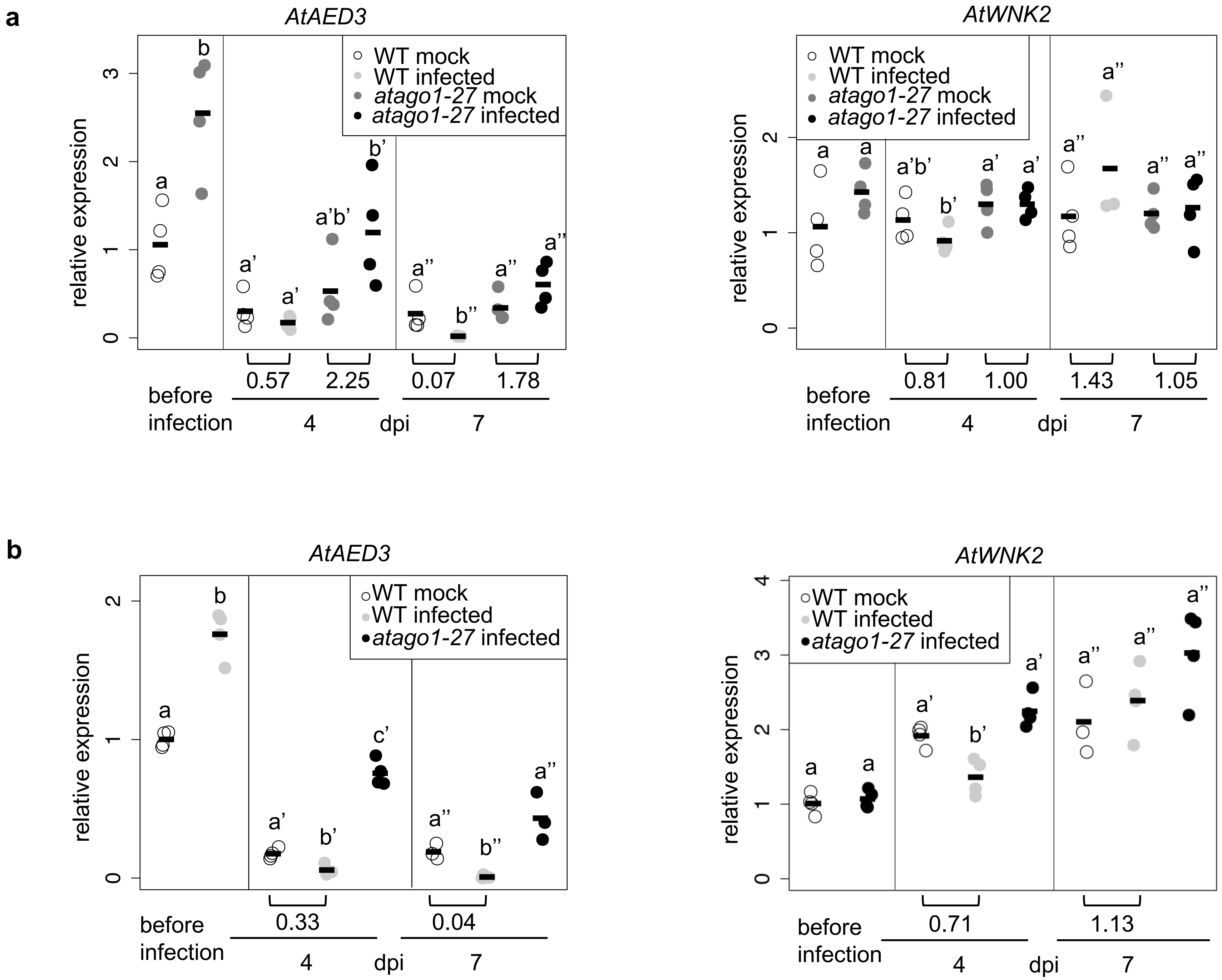
a
AtAED3
relative expression
WT mock
WT infected
atago1-27 mock
atago1-27 infected
0.57
2.25
0.07
1.78
before infection
4
dpi
7
AtWNK2
0.81
1.00
1.43
1.05
b
AtAED3
0.33
0.04
AtWNK2
0.71
1.13

Figure S4

**a**

```
AtAED3r     5' ACTGATGTTTATGGTACTCGA 3'
               ||0 ||||0|||0|   |  0|
HpasRNA90   3' CGATCACAAGTACTAGTATTT 5'
               |||| ||||||||0||  ||0|
AtAED3      5' GCTAATGTTCATGGTCCTAGA 3'

AtWNK2r     5' GAAACCCTGAAGAGCTGGAAA 3'
               | |  ||0|0 0|0 |00| |
HpasRNA2    3' CCTCAGGGCTCTTTAATTTCT 5'
               ||| |||0|||0|||||0|||
AtWNK2      5' GGAATCCTGAGGAATTAGAGA 3'
```

**b**

```
ATGGCCTCCTCAAGTCTCCATTTCTTCTTCTTCTTGACACTTCTCTTGCCATTCACTTTCACCACCGCCACACGTGACACATGTGCGACCGCCGCTCCCGATGGT
TCCGACGACCTCTCAATCATCCCCATTAACGCCAAATGCTCACCTTTCGCACCCACTCACGTCTCTGCCTCCGTGATAGACACGGTCCTTCACATGGCCTCCTCA
GACTCCCATCGACTCACCTACCTCTCAAGCCTCGTCGCCGGCAAACCAAAACCCACCTCCGTCCCCGTCGCCTCCGGTAATCAGCTCCACATCGGAAACTACGTG
                                                                                                      1/7       1/7
GTCCGAGCCAAACTCGGCACTCCTCCACAGCTAATGTTCATGGTCCTAGACACAAGTAACGACGCCGTTTGGCTCCCTTGCTCCGGCTGCTCCGGTTGTTCCAAC
                                                        1/7
GCCTCAACTTCTTTCAATACAAACTCCTCATCCACTTACTCAACCGTCTCTTGCTCCACCGCACAATGCACCCAAGCACGTGGCCTCACGTGCCCATCCTCCTCA
                                                1/7                                  1/7                    1/7
CCACAACCGTCCGTCTGCTCCTTCAACCAATCCTACGGCGGAGATTCATCCTTCTCTGCCAGTCTCGTCCAAGACACGCTAACGTTAGCCCCTGACGTCATCCCA
AATTTCTCCTTCGGCTGCATCAACTCCGCCTCAGGCAATTCTCTACCACCGCAAGGACTAATGGGCCTAGGACGCGGGCCTATGTCATTGGTCTCTCAAACTACG
TCGCTTTACTCAGGAGTGTTCTCATACTGCCTCCCTAGCTTCAGGTCCTTCTACTTCTCCGGGTCGTTGAAACTGGGTCTCTTGGGTCAACCCAAATCCATCAGA
TACACTCCACTCCTCCGTAACCCTCGCCGTCCATCACTTTACTATGTGAATCTCACCGGAGTGAGTGTCGGTTCCGTCCAGGTACCCGTTGACCCAGTGTATTTG
           1/7
ACGTTTGACGCAAATTCCGGCGCCGGTACGATCATCGATTCCGGTACTGTTATCACCCGTTTTGCTCAACCCGTGTACGAAGCCATTAGAGACGAGTTTAGGAAG
CAGGTGAATGTGTCGTCATTCTCAACATTGGGAGCGTTTGACACGTGTTTCTCAGCGGATAACGAGAATGTGGCGCCAAAGATAACGCTGCACATGACGTCACTC
GACTTGAAGCTGCCGATGGAGAATACGCTTATCCACAGCAGCGCGGGAACGCTGACGTGTCTATCCATGGCCGGGATACGGCAGAACGCAAACGCTGTTCTAAAC
GTGATCGCGAATCTCCAACAGCAGAACTTAAGGATCTTGTTTGACGTTCCTAATTCTCGCATAGGAATTGCTCCTGAGCCCTGCAACTAA
```

**Figure S5**

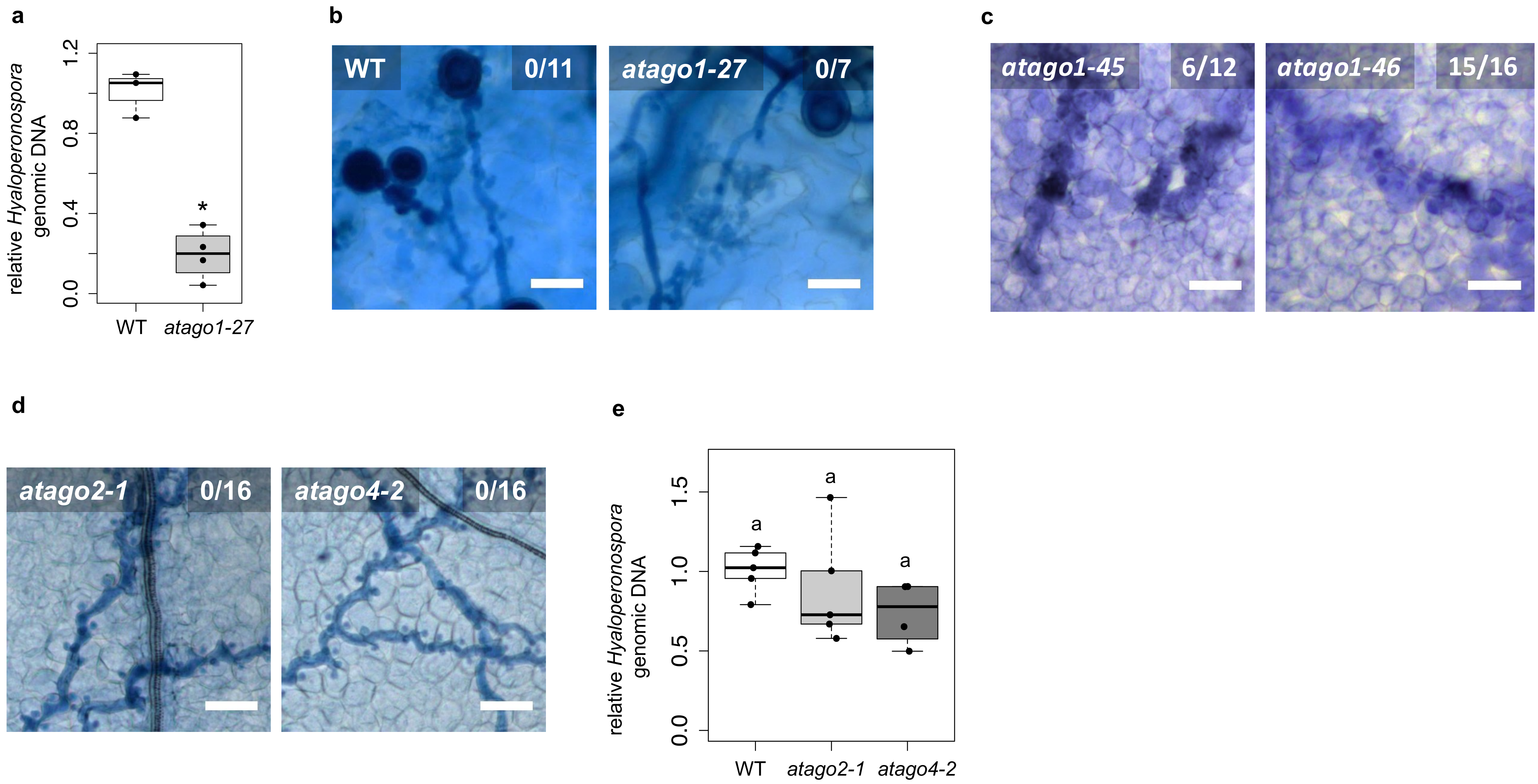
a
relative Hyaloperonospora genomic DNA
0.0
0.4
0.8
1.2
*
WT
atago1-27
b
WT
0/11
atago1-27
0/7
c
atago1-45
6/12
atago1-46
15/16
d
atago2-1
0/16
atago4-2
0/16
e
relative Hyaloperonospora genomic DNA
0.0
0.5
1.0
1.5
a
a
a
WT
atago2-1
atago4-2

**Figure S6**

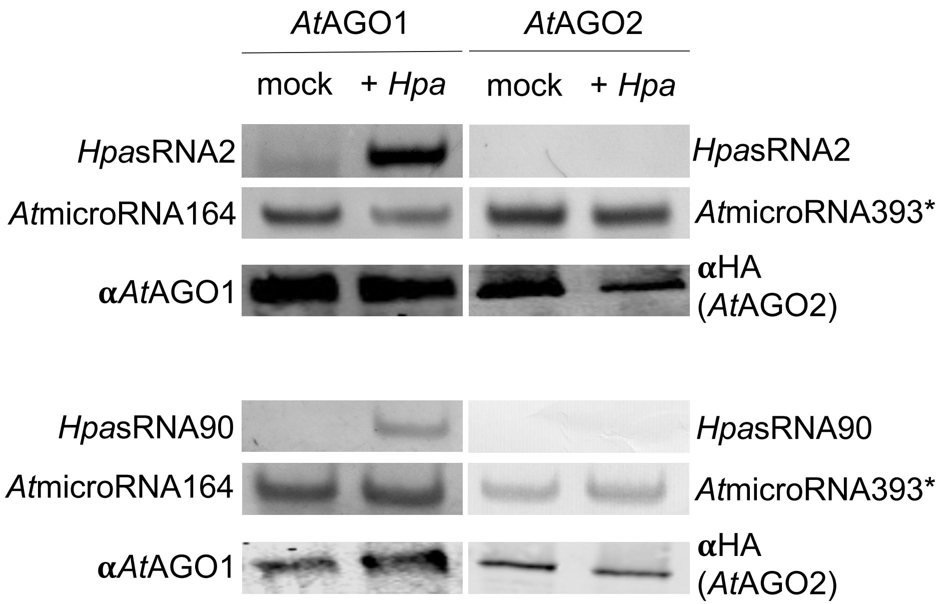
*At*AGO1
*At*AGO2
mock
+ *Hpa*
mock
+ *Hpa*
*Hpa*sRNA2
*At*microRNA164
α*At*AGO1
*Hpa*sRNA2
*At*microRNA393*
αHA (*At*AGO2)
*Hpa*sRNA90
*At*microRNA164
α*At*AGO1
*Hpa*sRNA90
*At*microRNA393*
αHA (*At*AGO2)

**Figure S7**

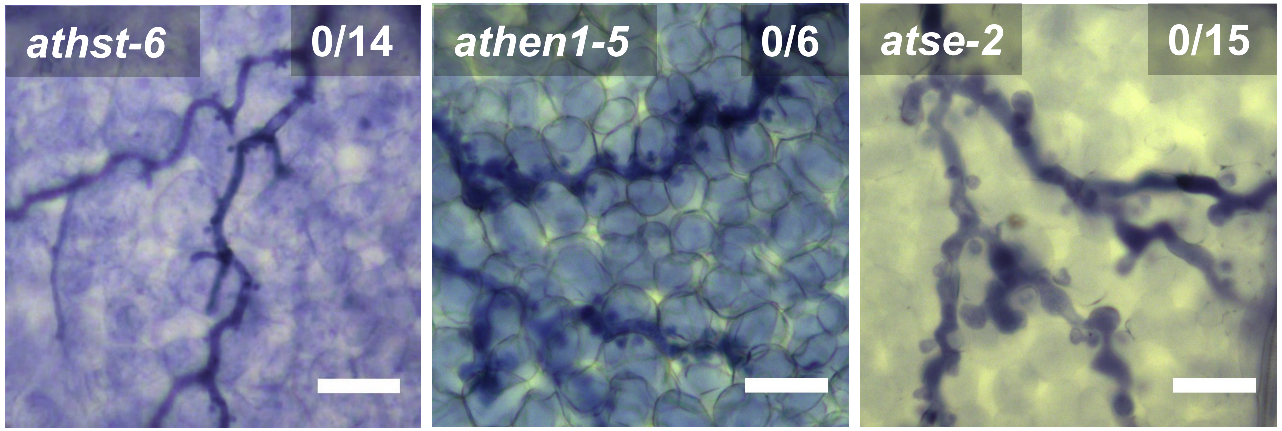


**Figure S8**

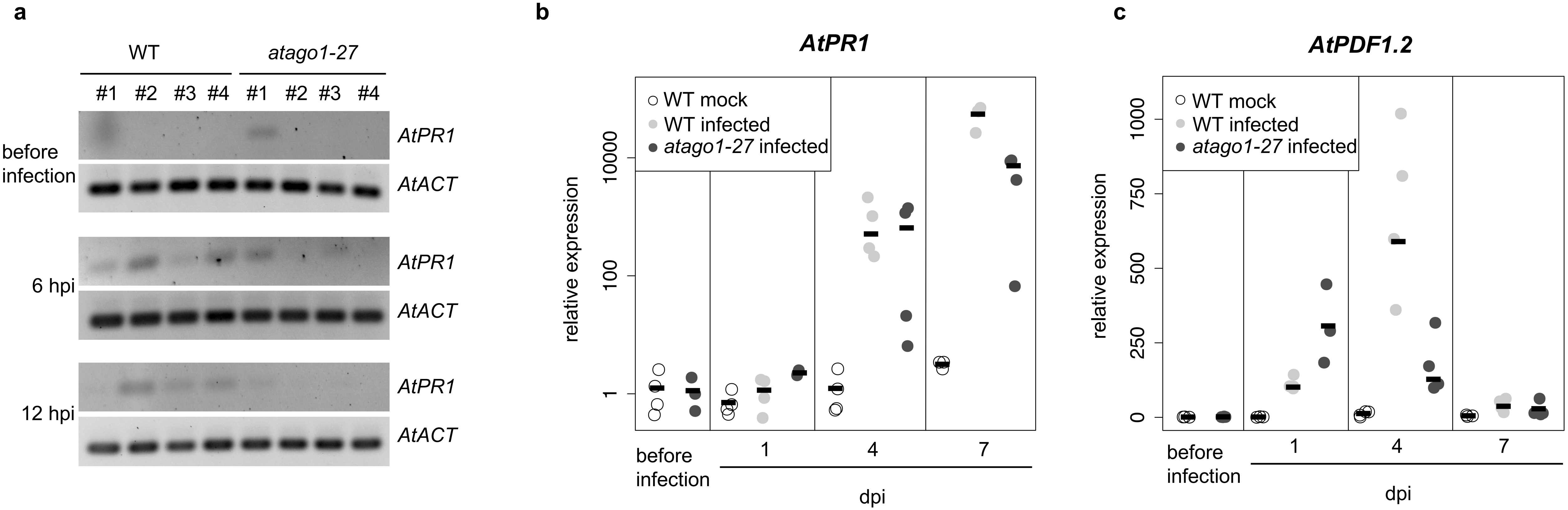
a
WT
atago1-27
#1
#2
#3
#4
#1
#2
#3
#4
before infection
6 hpi
12 hpi
AtPR1
AtACT
AtPR1
AtACT
AtPR1
AtACT
b
AtPR1
WT mock
WT infected
atago1-27 infected
relative expression
10000
100
1
before infection
1
4
7
dpi
c
AtPDF1.2
WT mock
WT infected
atago1-27 infected
relative expression
1000
750
500
250
0
before infection
1
4
7
dpi

**Figure S9**

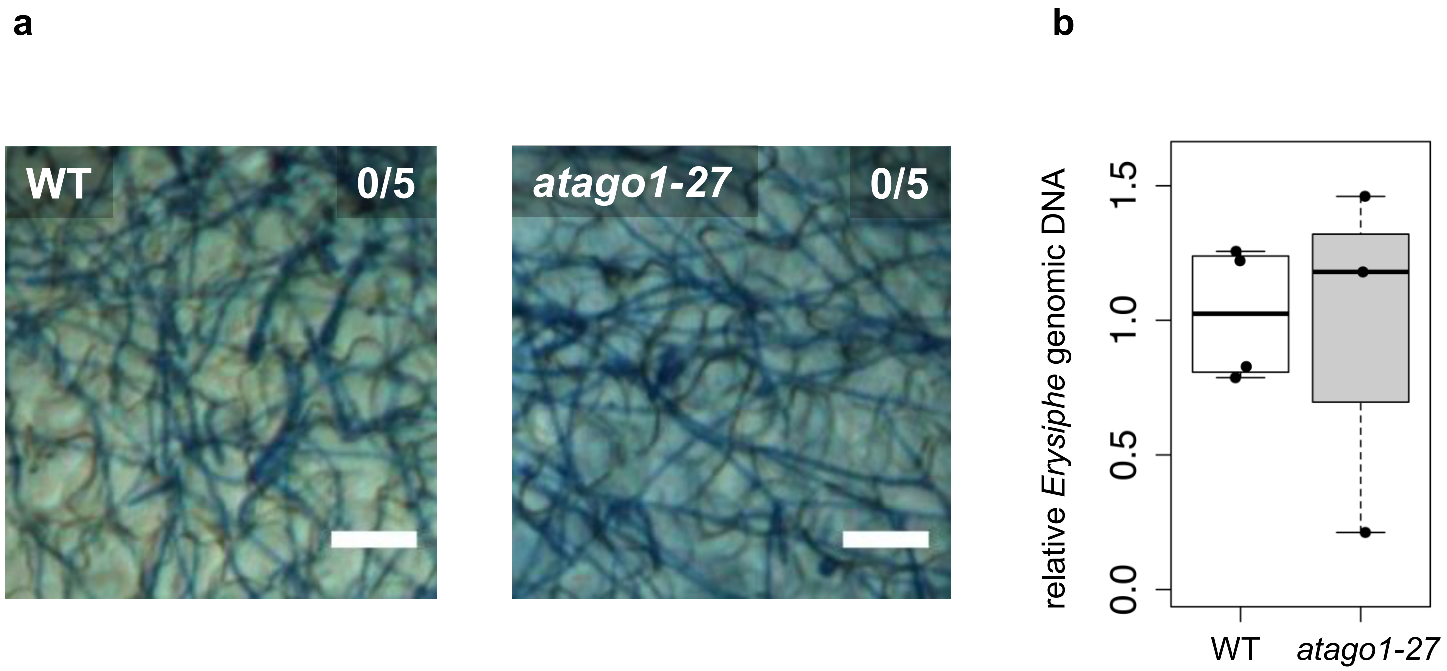
a
WT
0/5
atago1-27
0/5
b
relative Erysiphe genomic DNA
0.0
0.5
1.0
1.5
WT
atago1-27

**Figure S10**

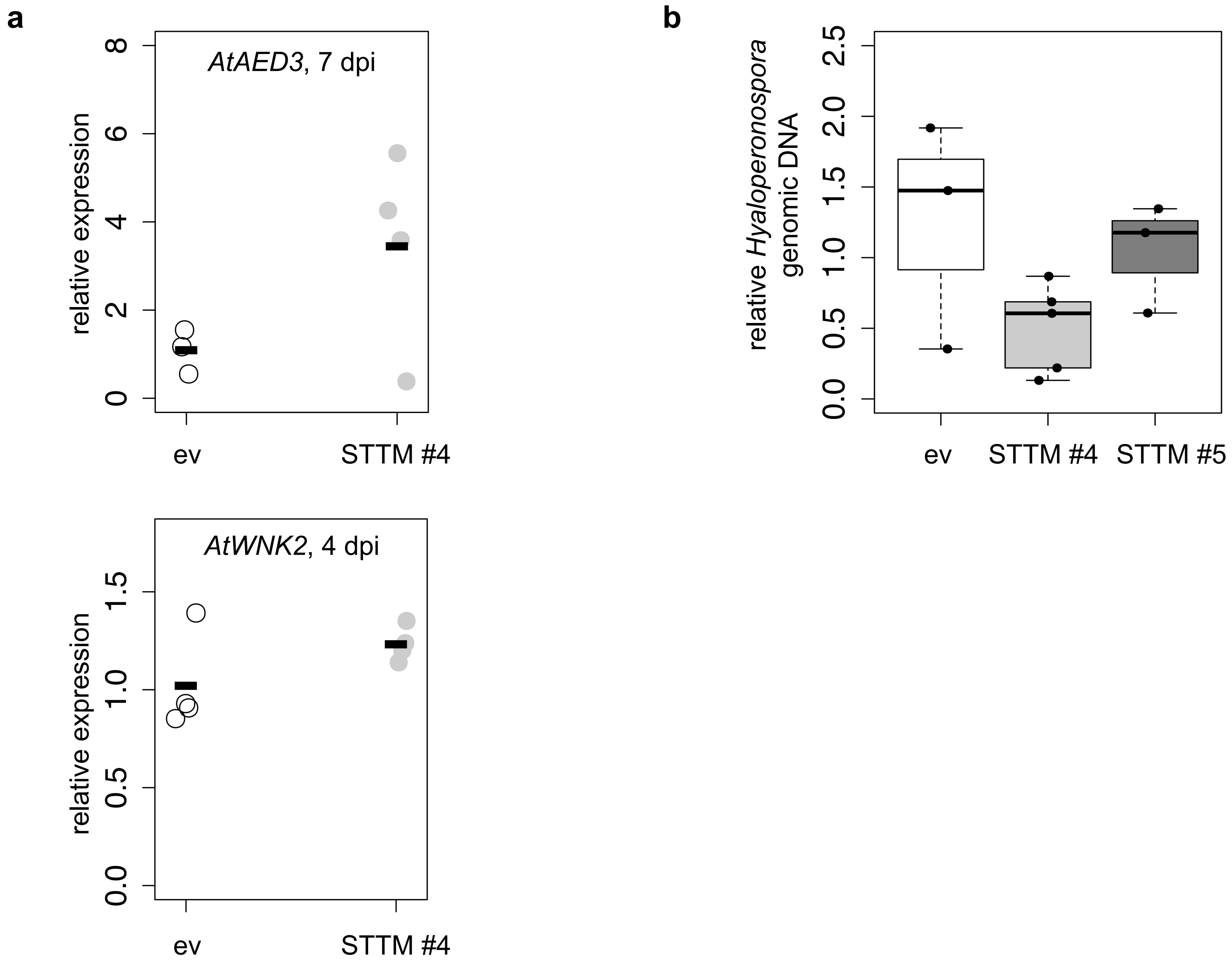
a
AtAED3, 7 dpi
relative expression
0
2
4
6
8
ev
STTM #4
AtWNK2, 4 dpi
relative expression
0.0
0.5
1.0
1.5
ev
STTM #4
b
relative Hyaloperonospora genomic DNA
0.0
0.5
1.0
1.5
2.0
2.5
ev
STTM #4
STTM #5

**Figure S11**

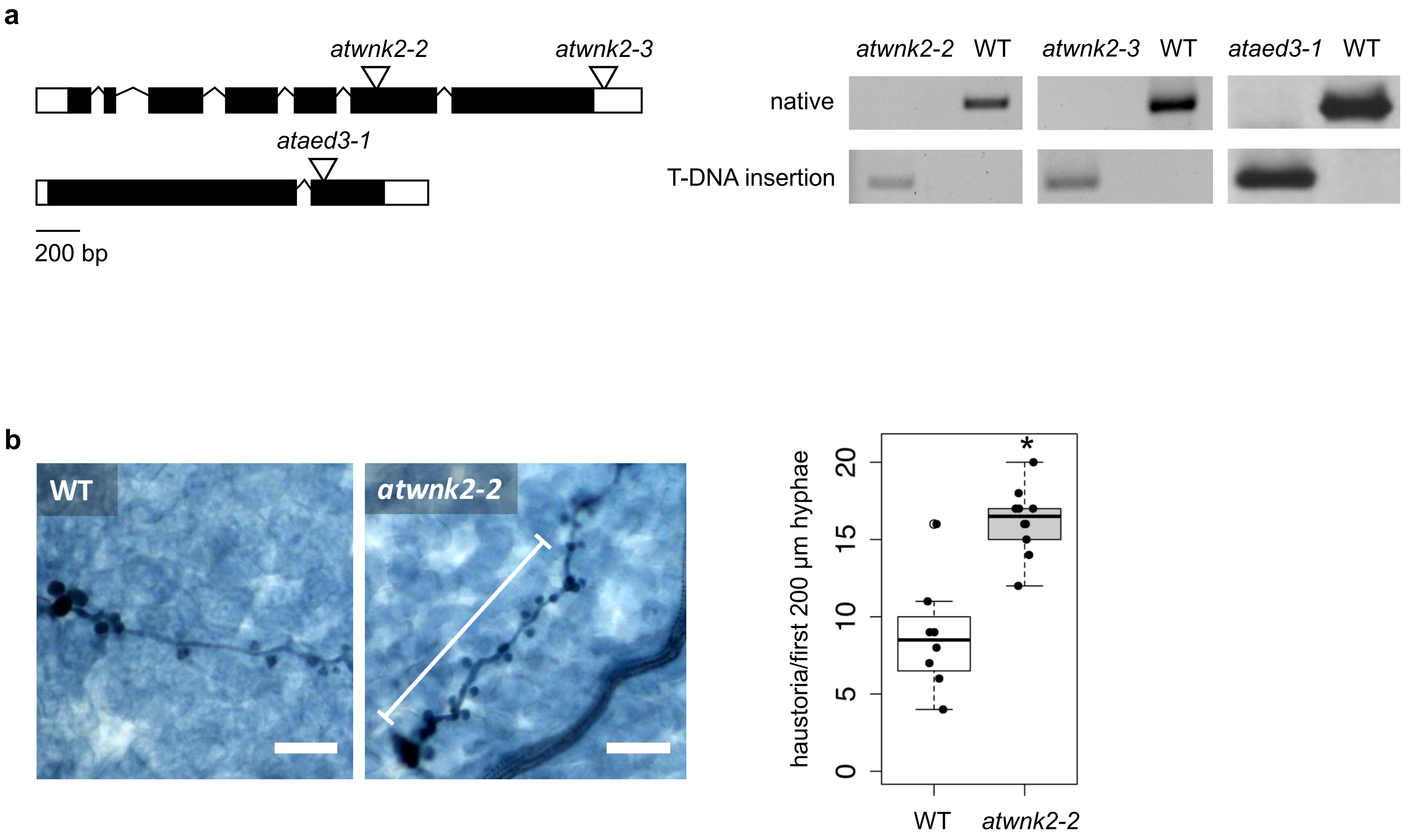
a
atwnk2-2
atwnk2-3
ataed3-1
200 bp
atwnk2-2 WT
atwnk2-3 WT
ataed3-1 WT
native
T-DNA insertion
b
WT
atwnk2-2
haustoria/first 200 µm hyphae
0
5
10
15
20
*
WT
atwnk2-2

**Figure S12**

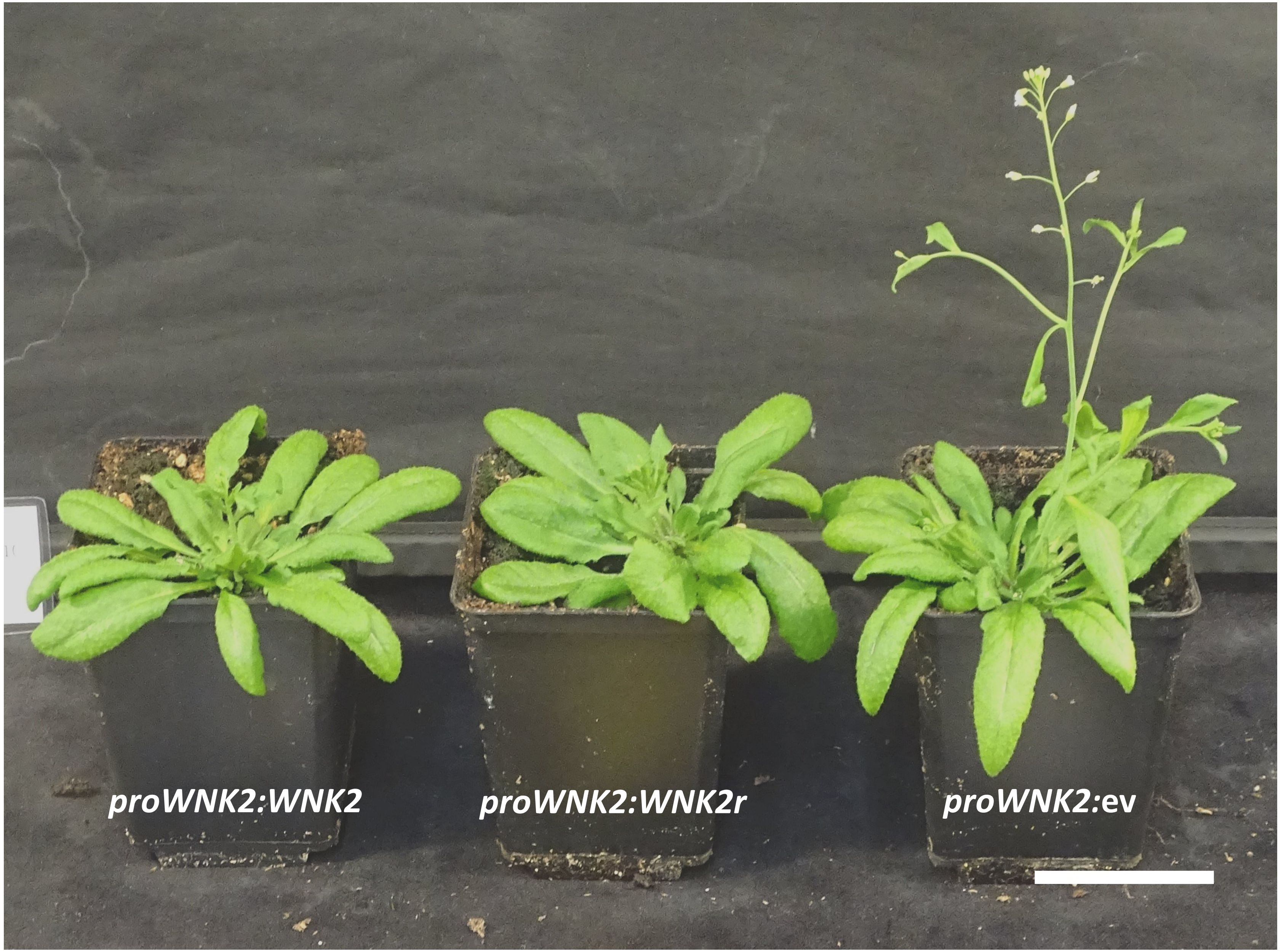


**Figure S13**

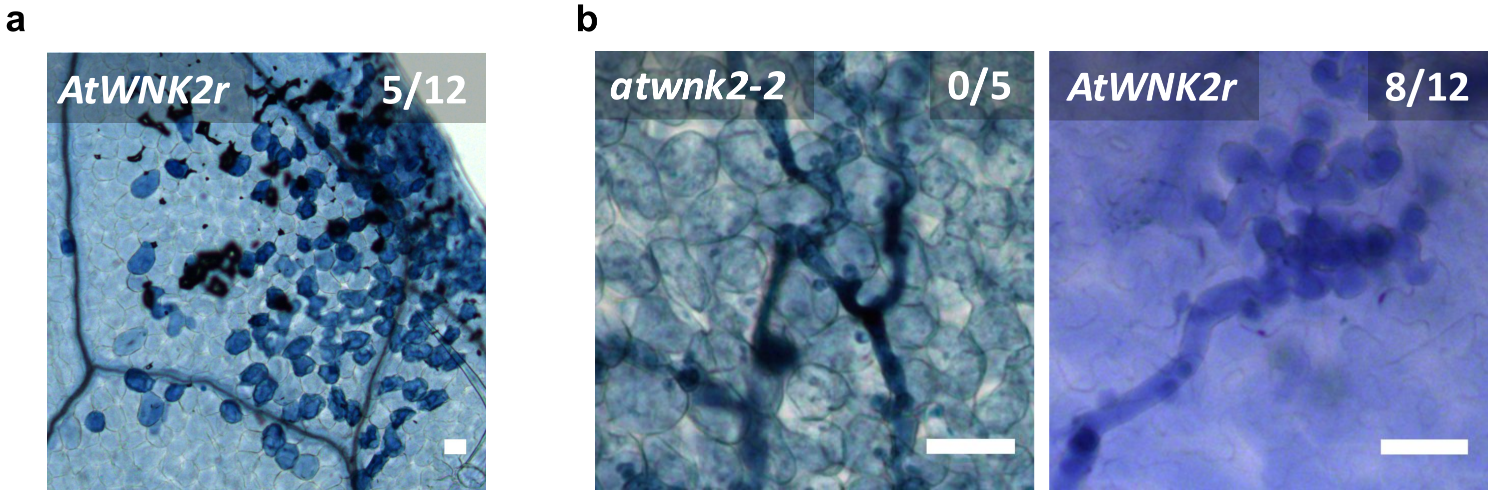


**Figure S14**

**a**

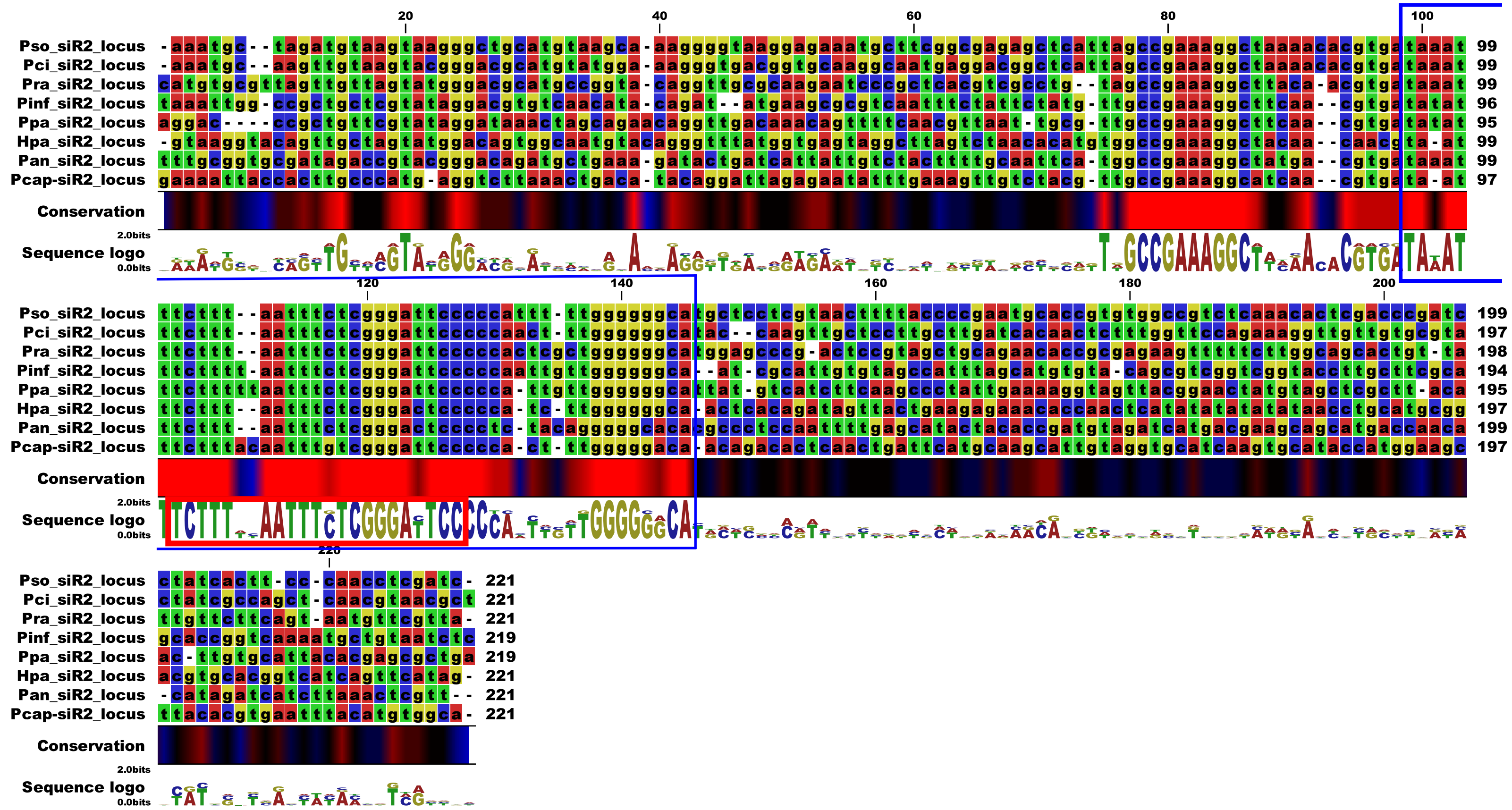


**b**

```
HpasRNA2      5’  TCTTTAATTTCTCGGGACTCC 3’
                  |||0|||||0|||0||| |||
AtWNK2        3’  AGAGATTAAGGAGTCCTAAGG 5’

PcapsRNA2     5’  TTTACAATTTGTCGGGATTCC 3’
                  |0|  ||||00||0|||||||
AtWNK2        3’  AGAGATTAAGGAGTCCTAAGG 5’

PsosRNA2      5’  TCTTTAATTTCTCGGGATTCC 3’
                  |||0 |   |0 ||0|||||||
GmWNK2        3’  AGAGTTCTAGAAGTCCTAAGA 5’

PinfsRNA2     5’  TCTTTTAATTTCTCGGGATTCC 3’
                  |||0   |   |0 ||0|||0|
StWNK2        3’  AGAGT-TCTAGAAGTCCTGAAA 5’

PpasRNA2      5’  TCTTTTTAATTTCTCGGGATTCC 3’
                  |||0    |   |0 ||0|||0|
NtWNK2        3’  AGAG--TTCTAGAAGTCCTGAAA 5’
```

Figure S15
